# Controlled human infection with *Plasmodium falciparum-*infected mosquito bites elicits antibodies against mosquito salivary protein SG1L3

**DOI:** 10.64898/2026.03.19.713001

**Authors:** Carolina M. Andrade, Renate C. van Daalen, Amanda Fabra-García, Sanne Grievink, Geert-Jan van Gemert, Karina Teelen, Svenja Hester, Rianne Stoter, Marga van de Vegte-Bolmer, Chris Drakeley, Alfred B. Tiono, Robert W. Sauerwein, Teun Bousema, Matthijs M. Jore

## Abstract

Human malaria infections begin with the injection of Plasmodium sporozoites via mosquito saliva. Whole sporozoite immunizations have been used as a model to study immune responses to malaria parasites, having culminated in circumsporozoite protein (CSP)-targeting vaccines and monoclonal antibodies (mAbs). However, antibody responses targeting non-CSP antigens on the sporozoite surface remain poorly characterized. Here, we isolated single B cells from a human volunteer immunized by *Plasmodium falciparum-*infected mosquito bites, who had acquired non-CSP-specific antibodies that recognize sporozoites. We identified two mAbs that recognize the surface of *P. falciparum* sporozoites, but do not bind to CSP. Using immunoprecipitation followed by mass-spectrometry, we found that the target of these mAbs is not a *P. falciparum* protein but the mosquito salivary protein SG1L3. We observed that recombinant SG1L3 binds to *P. falciparum* sporozoites. However, the SG1L3-specific mAbs and SG1L3-specific polyclonal antibodies from this volunteer, as well as polyclonal antibodies raised against recombinant SG1L3 in rabbits, fail to block liver stage infection *in vitro*, making this an unlikely target for functional antibodies. We observed that inhabitants from an area with intense *Anopheles* exposure in Burkina Faso can have antibodies against SG1L3, and that antibody titers increase with age. In conclusion, we identified the first human mAbs against a mosquito saliva protein that binds to the surface of sporozoites. Future work should assess whether naturally acquired antibodies against this protein may be used as a serological marker of mosquito exposure.

## Introduction

Malaria control efforts have made significant progress, with more than two billion cases and 10 million deaths averted in the last two decades (World Health Organization 2025). However, elimination efforts have recently stalled. In 2024, malaria was responsible for 282 million cases and 610,000 deaths (World Health Organization 2025). The majority of these deaths occurred in African children under the age of five, where *P. falciparum* dominates infections.

*P. falciparum* has a complex life-cycle, that alternates between the mosquito vector and the human host. Human infection starts when an infected female *Anopheles* mosquito takes a bloodmeal and inoculates parasites mixed with mosquito saliva in the human skin. The parasites then travel to the liver, where they cross the liver sinusoids and traverse several hepatocytes before invading a final hepatocyte. During its development inside a hepatocyte, a single *P. falciparum* replicates and forms thousands of new parasites that emerge from the hepatocyte. These parasites initiate the erythrocytic stage, during which they infect and replicate inside red blood cells, causing malaria symptoms and disease. The pre-erythrocytic stage of the infection constitutes an attractive target for interventions due to its clinically silent nature, offering the possibility of preventing malaria disease and transmission (Prudencio, Rodriguez et al. 2006).

Early studies from Nussenzweig and colleagues have shown that radiation attenuated sporozoites are able to induce protective immunity in mice (Nussenzweig, Vanderberg et al. 1967). Subsequently, Clyde et al. showed that high levels of immunity are achievable in naïve human volunteers following immunization with irradiated *P. falciparum* sporozoites (Clyde, McCarthy et al. 1973). These seminal studies demonstrated that it is possible to target *Plasmodium* sporozoites to prevent infection. The main target of the antibodies generated by these immunizations was later identified to be circumsporozoite protein (CSP), an immunodominant protein present at the surface of the sporozoite (Nussenzweig and Nussenzweig 1984). CSP is essential for sporozoite formation and development in the mosquito, and infection of the human host and invasion of hepatocytes (Singer, Kanatani et al. 2024).

There are currently two licensed vaccines against malaria, RTS,S/AS01_B_ and R21/Matrix-M. Both vaccines are recommended by the World Health Organization for prevention of malaria in children living in endemic regions (World Health Organization 2021, World Health Organization 2023). Both RTS,S and R21 vaccines target the *Plasmodium falciparum*’s CSP. Although these vaccines are likely to avert thousands of malaria deaths, their efficacy is incomplete and depends on annual boosters (RTS;S Clinical Trials Partnership 2015, Datoo, Dicko et al. 2024), underscoring the need to identify additional vaccine targets.

Natural exposure or experimental immunizations can induce antibodies against sporozoite surface antigens other than CSP (Doolan, Southwood et al. 2003, John, Moormann et al. 2005, Doolan, Mu et al. 2008, Felgner, Roestenberg et al. 2013, Nahrendorf, Scholzen et al. 2014, Obiero, Campo et al. 2019). Of these, Thrombospondin-Related Anonymous Protein (TRAP), Liver Stage Antigens 1 and 3 (LSA1, LSA 3), Cell Traversal protein for Ookinetes and Sporozoites (CelTOS), Apical Membrane Antigen 1 (AMA1) and Sporozoite Threonine and Asparagine Rich Protein (STARP) have been examined most extensively as potential vaccine targets in preclinical and clinical trials, (World Health Organization 2025) but showed little to no efficacy in preventing infection.

Recently, isolation of single B cells followed by recombinant monoclonal antibody (mAb) expression has enabled identification of functional mAbs from volunteers naturally or experimentally exposed to malaria causing parasites. Using this approach, several studies have identified potent mAbs that target CSP, and that are being studied as anti-infection prophylaxis (Oyen, Torres et al. 2017, Kisalu, Idris et al. 2018, Wang, Pereira et al. 2020, Williams, Guerrero et al. 2024). To this date, human mAbs that target non-CSP targets on the sporozoite surface have not yet been identified.

Here, we aimed to unravel new targets on the sporozoite surface that are recognized by antibodies from human volunteers that were immunized with whole sporozoites. To this end, we used an untargeted B-cell sorting approach, followed by recombinant mAb production, immunoprecipitation and mass-spectrometry for target identification and validation. We identified two mAbs that target the *Anopheles stephensi* mosquito salivary protein SG1L3 bound to the sporozoite surface but found that mono- and polyclonal antibodies against SG1L3 did not reduce liver stage infection *in vitro.* To our knowledge, our study is the first to report the discovery of human mAbs against non-CSP targets on the sporozoite surface.

## Results

### Target agnostic B-cell sorting identifies two unique non-CSP monoclonal antibodies

We previously immunized 12 malaria-naïve volunteers, receiving chemoprophylaxis, with *Plasmodium falciparum* NF54 sporozoites by *Anopheles stephensi* mosquito bite (Walk, Reuling et al. 2017) (Sup. Fig. 1a). We observed that most volunteers developed polyclonal antibodies that block *P. falciparum* sporozoite invasion of HC-04 liver cells *in vitro* and liver infection in a humanized mouse model (Fabra-Garcia, Yang et al. 2022) (Sup. Fig. 1b). Interestingly, after depletion of CSP-specific IgGs, polyclonal antibodies from several volunteers retained strong activity suggesting the presence of functional non-CSP targets on the sporozoite surface. These CSP-depleted polyclonal antibodies further retained recognition of *P. falciparum* sporozoites surface confirming the presence of non-CSP targets at their surface.

**Figure 1:**
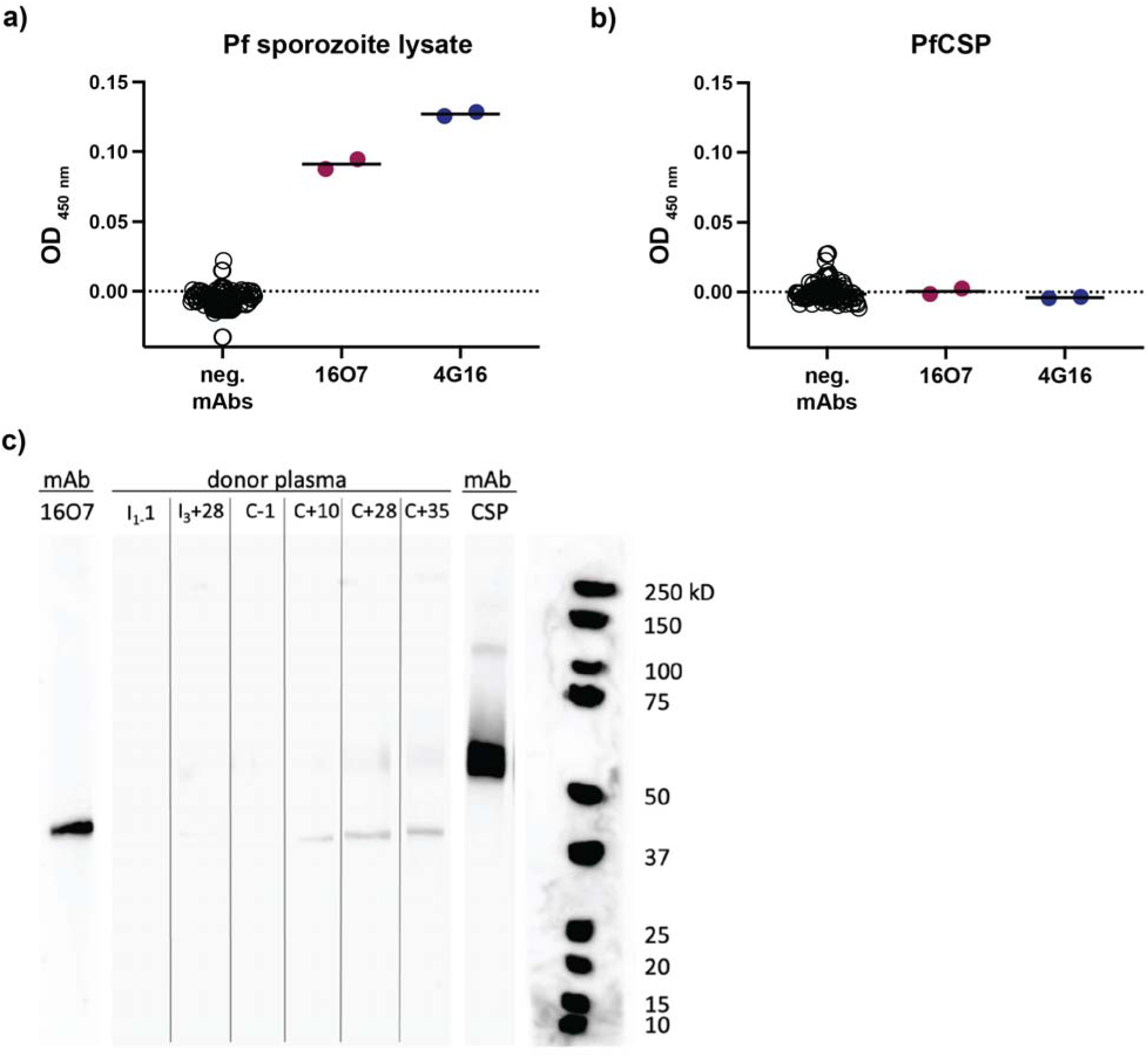
Target agnostic B-cell sorting identifies two non-CSP monoclonal antibodies **a)** Binding of mAbs in *P. falciparum* NF54 sporozoite lysate ELISA and **b)** binding of mAbs in recombinant full length CSP ELISA. Supernatants of HEK293T cells expressing mAbs were tested at 1:5 dilution. mAbs that showed no reactivity in both ELISAs are grouped as negative mAbs (each ran as duplicates). Data represents OD_450nm_ minus average of blank. Bars for 16O7 and 4G16 represent mean from two technical replicates. Anti-CSP mAb was included as positive control (not shown). n= 55 mAbs (including 16O7 and 4G16 mAbs). **c)** Western blot strips with *P. falciparum* NF54 sporozoite lysate, incubated with monoclonal antibody 16O7 (1 µg/ml), CSP monoclonal antibody CIS43 (0.08 µg/ml) or plasma (1:5 dilution) from the volunteer at different time-points. Time-points: I1-1 inclusion plasma; I3+28, 28 days post third immunization; C-1, one day prior to challenge; C+10, 28 or 35: 10-, 28- or 35-days post challenge (see supplementary figure for immunization regimen).

To identify the target(s) of these non-CSP-specific antibodies, we selected the volunteer with the strongest non-CSP activity in both assays for B-cell analysis (Sup Fig 1b; volunteer 1 in Fabra-Garcia, Yang et al 2022). We performed untargeted B-cell sorting, and sorted 5071 single B cells (CD19^+^ IgM^-^ IgA^-^ IgD^-^, gating strategy in Sup. Fig 1c). B cells were then stimulated for two weeks, and secreted IgGs were screened for reactivity against *P. falciparum* sporozoite lysate (Sup. Fig. 1d). We selected 264 samples (∼5%) with the highest relative reactivity to perform nested PCR on their B cell receptor for both heavy and light chains. We then cloned these into IgG1 expression vectors, and transfected HEK293T cells to recombinantly produce monoclonal antibodies (mAbs). Of the selected samples, 55 samples were successfully cloned and yielded detectable IgG1 in the supernatant. Of these, 53 mAbs showed no reactivity against recombinant CSP or sporozoite lysate. Two mAbs, 16O7 and 4G16 (Sup. Table 1 for VDJ gene usage), were reactive against *P. falciparum* sporozoite lysate by ELISA (Fig. 1a-b), but not against recombinant CSP and were examined further. These mAbs were isolated from B cells whose supernatants were amongst the five highest reactivities against sporozoite lysate (Sup. Fig. 1d).

We performed a western blot with *P. falciparum* sporozoite lysate and observed that mAb 16O7 recognizes a protein of approximately 40-45 kDa, distinct from CSP that runs above 50 kDa (Fig. 1c). In parallel, we conducted a western blot with volunteers’ plasma and found that the volunteer had mainly developed antibodies against a protein of similar size as that recognized by mAb 16O7, as well as antibodies against a protein of similar size as CSP. Of note, we did not detect any signal with the mAb 4G16 by western blot, possibly due to the recognition of a conformational epitope that is sensitive to the denaturing SDS conditions.

Taken together, we identified two mAbs from a volunteer with strong non-CSP-specific anti-sporozoite activity, one of which recognizes a protein of 40-45 kDa.

### Non-CSP mAbs recognize a salivary protein deposited at the surface of the sporozoite

As sporozoite lysate is generated from dissected *P. falciparum*-infected *Anopheles* mosquitoes’ salivary glands, we also tested recognition of lysate from uninfected mosquito salivary glands and found that the 40-45 kDa protein recognized by mAb 16O7 is also present in saliva of uninfected *Anopheles stephensi* mosquitoes (Fig. 2a).

**Figure 2:**
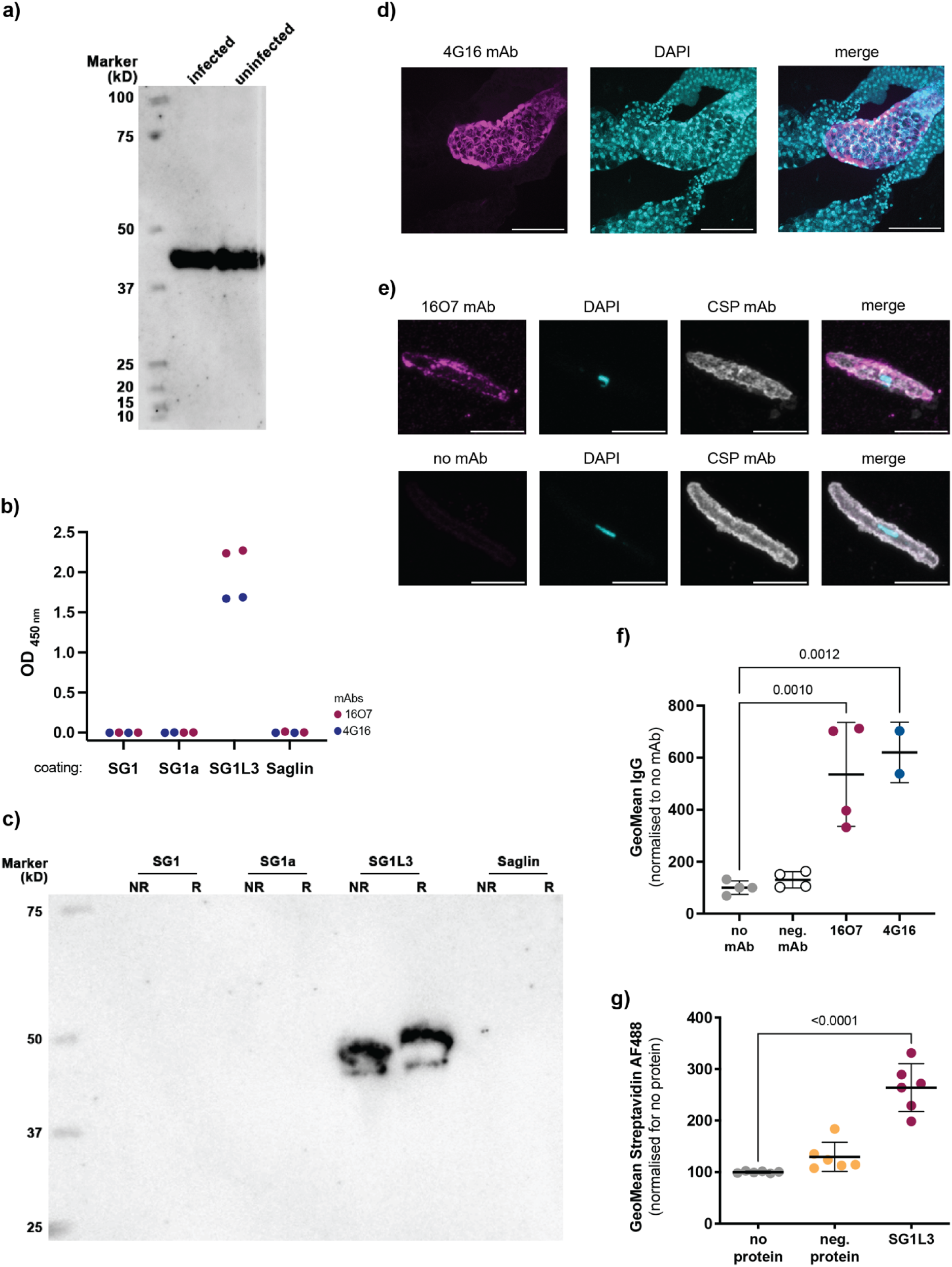
Non-CSP mAbs recognize a salivary protein deposited at the surface of *P. falciparum* sporozoites **a)** Western blot with 16O7 mAb, with either *Pf* NF54-infected or uninfected *Anopheles stephensi* salivary gland lysate. **b)** ELISA with mAbs 16O7 (magenta) and 4G16 (blue), with different recombinant SG1 family proteins used for coating. Each mAb was tested in duplicate per condition. **c)** Western blot with 16O7 mAb, with several SG1 family recombinant proteins under reducing (R) and non-reducing (NR) conditions. **d)** Immunofluorescence of *Anopheles stephensi* mosquito salivary gland with 4G16 mAb (magenta). Cyan, DAPI. Picture shows maximum projection intensity. Scale bar 100 µm. **e)** *P. falciparum* sporozoite staining with 16O7 mAb (top) or no primary antibody (bottom). Magenta, 16O7 mAb; Cyan, DAPI; Gray, CSP mAb. Scale bar 5 µm. **f)** Sporozoite binding of mAbs to sporozoites by flow cytometry. GeoMean of IgG+ sporozoites, normalized to no mAb control. mAb TB31F that recognizes gametes was used as negative control (neg mAb). Two independent experiments with two technical replicates each, except 4G16 (only one experiment). **g)** Recombinant SG1L3 protein binding to *P. falciparum* sporozoites by flow cytometry. Binding was detected using a biotinylated anti-c-tag antibody. Values are GeoMean of Streptavidin Alexa Fluor (AF) 488, normalised to the no protein control. Recombinant Pfs230D10 containing a C-tag was used as negative control (neg. protein). Three independent experiments with two technical replicates f-g) Data were analysed using one-way ANOVA with Dunnet’s multiple comparisons test; Bars represent mean ± SD (standard deviation). p>0.05 (non-significant) values are not shown.

In order to identify the target of mAb 16O7, we performed immunoprecipitation with uninfected salivary gland extract, followed by liquid-chromatography mass-spectrometry. Mass-spectrometry identified SG1L3 as the protein with the highest intensity value, and further identified several other members from the SG1 family (Sup. Table 2). We recombinantly expressed salivary protein SG1L3, and three other *A. stephensi* SG1 family members that share the highest amino acid identity to SG1L3 (SG1, SG1a and saglin) (Sup. Fig. 2). Both ELISA (Fig. 2b) and western blot (Fig. 2c) showed that mAbs 16O7 and 4G16 recognized SG1L3, but not the other SG1 family proteins, indicating they recognize SG1L3-specific epitopes. Using mAb 4G16 in immunofluorescence microscopy, we observed that salivary protein SG1L3 is present in the median lobe of *A. stephensi* salivary glands (Fig. 2d).

We then explored if SG1L3 is present at the surface of *P. falciparum* sporozoites. We performed immunofluorescence microscopy and observed that mAb 16O7 binds to the surface of *P. falciparum* sporozoites (Fig. 2e), whereas no signal was observed in the absence of mAbs. Additionally, we evaluated binding of mAbs 1607 and 4G16 by flow cytometry and found that both mAbs recognized *P. falciparum* sporozoites (Fig. 2f). Together, these results suggest that SG1L3 salivary protein is deposited at the surface of *P. falciparum* sporozoites. Of note, we could not detect any recognition of gametes by our mAbs (Sup. Fig. 3 a), whereas a clear signal was present when transmission-blocking mAb TB31F was used as a positive control in this immunofluorescence assay.

**Figure 3:**
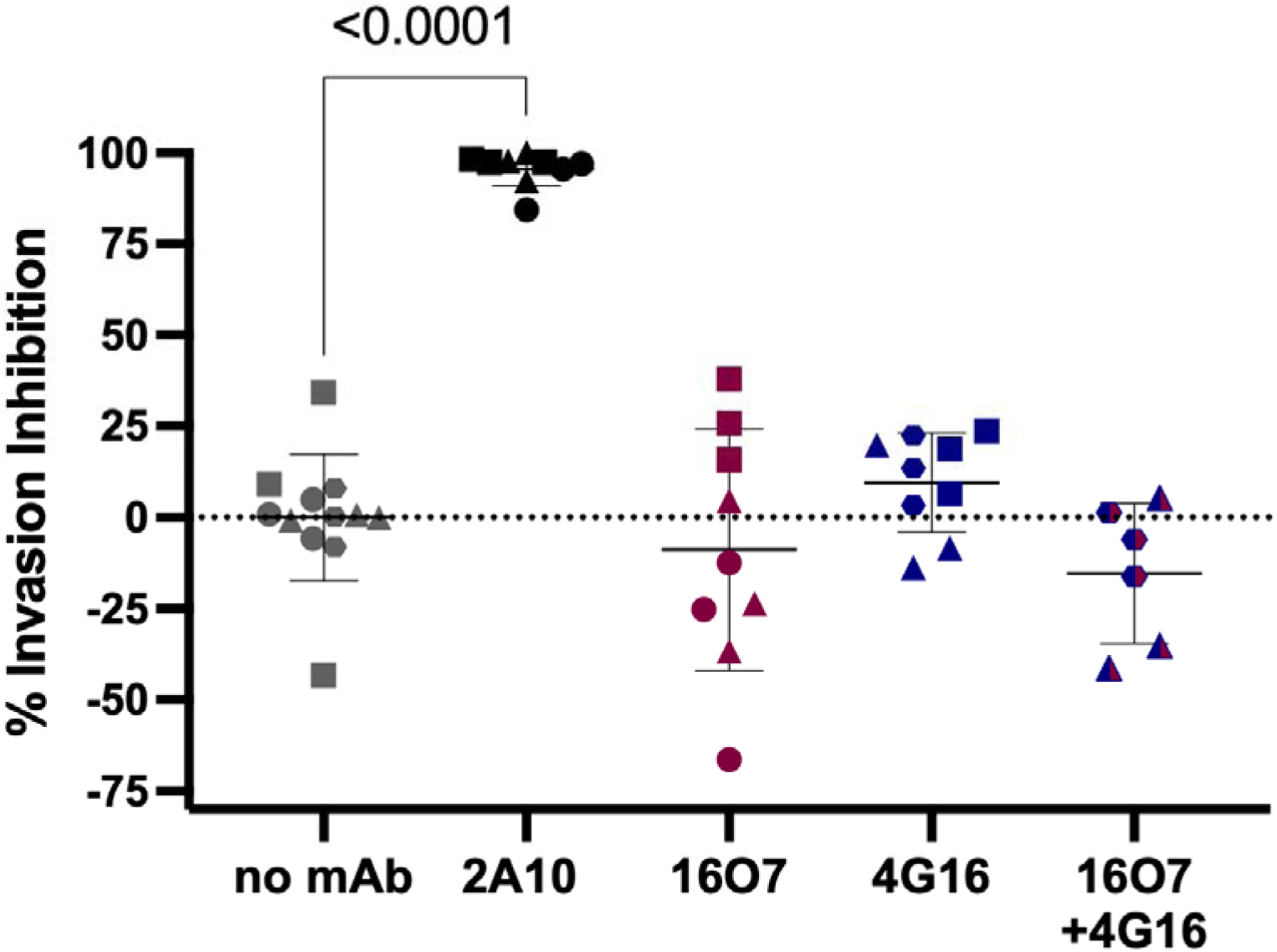
SG1L3 mAbs show no invasion inhibition of *P. falciparum* sporozoites Invasion inhibition of *P. falciparum* sporozoites of HC-04 cells when incubated with 100 µg/ml mAb, or a combination of both monoclonal antibodies (50µg/ml 16O7 + 50µg/ml 4G16). Inhibition was calculated as the reduction in number of infected HC-04 cells and normalized against the no mAb control. 2A10 (10 µg/mL) is a CSP-specific antibody used as positive control. Data were analyzed using one-way ANOVA with Dunnett’s multiple comparisons test compared to the no mAb control; p>0.05 values are not shown. mAbs 2A10, 16O7 and 4G16 were tested in three independent experiments with three technical replicates each, 16O7+4G16 was tested in two independent experiments with three technical replicates each. Different shapes indicate independent experiments. Error bars represent mean ± SD (standard deviation).

To confirm that SG1L3 is binding to the sporozoite surface, we performed a binding assay with recombinant C-tagged SG1L3 or an irrelevant C-tagged protein as negative control. Using an anti C-tag antibody, we measured the fluorescence at the surface of sporozoites by flow cytometry. We observed that the SG1L3 protein binds to the surface of *P. falciparum* sporozoites (Fig. 2g), whereas the negative control protein does not. To test if the observed interaction of SG1L3 was specific to sporozoites, we also performed a binding assay with gametes. We could not detect any signal above baseline when gametes were incubated with recombinant SG1L3 protein (Sup. Fig. 3 b), whereas a positive control protein showed strong signal.

Our data shows that both mAbs specifically recognize *A. stephensi* salivary protein SG1L3, and that SG1L3 binds to the surface of *P. falciparum* sporozoites.

### SG1L3 mAbs do not inhibit hepatocyte invasion

mAbs 16O7 and 4G16 were obtained from a volunteer that had polyclonal antibodies that blocked *in vitro* invasion of HC-04 hepatocyte cells. Therefore, we tested whether these mAbs were functional in the same assay. *P. falciparum* sporozoites were incubated with 100 µg/ml mAbs prior to invasion, and hepatocyte invasion was measured by flow cytometry. We observed no invasion inhibition when *P. falciparum* sporozoites were incubated with mAb 16O7, mAb 4G16 or a combination of both (Fig. 3). As expected, positive control CSP mAb 2A10 showed strong inhibition of hepatocyte invasion.

### Polyclonal SG1L3 IgGs from immunized volunteer and immunized rabbits do not inhibit hepatocyte invasion and traversal

To assess if SG1L3 can elicit antibodies able to inhibit *P. falciparum* sporozoite infection *in vitro*, we used two approaches: i) Purification and depletion of SG1L3 polyclonal IgGs from the immunized volunteer and ii) rabbit immunization with recombinant SG1L3 salivary protein.

First, we covalently coupled recombinant SG1L3 protein to a column to purify and deplete SG1L3-specific IgGs from the volunteer’s polyclonal IgGs at 28 days post challenge (C+28) (Sup. Fig 1a). We confirmed successful depletion of SG1L3-specific IgGs to baseline levels (inclusion) by ELISA (Sup. Fig. 4 a). We also tested depletion for SG1L3-specific antibodies by western blot with *P. falciparum*-infected salivary gland extract and found that the reactivity with the 40-45 kDa protein disappeared (Sup. Fig. 4b). This result demonstrates that SG1L3 is one of the two major antibody targets in this volunteer (the other being CSP).

**Figure 4:**
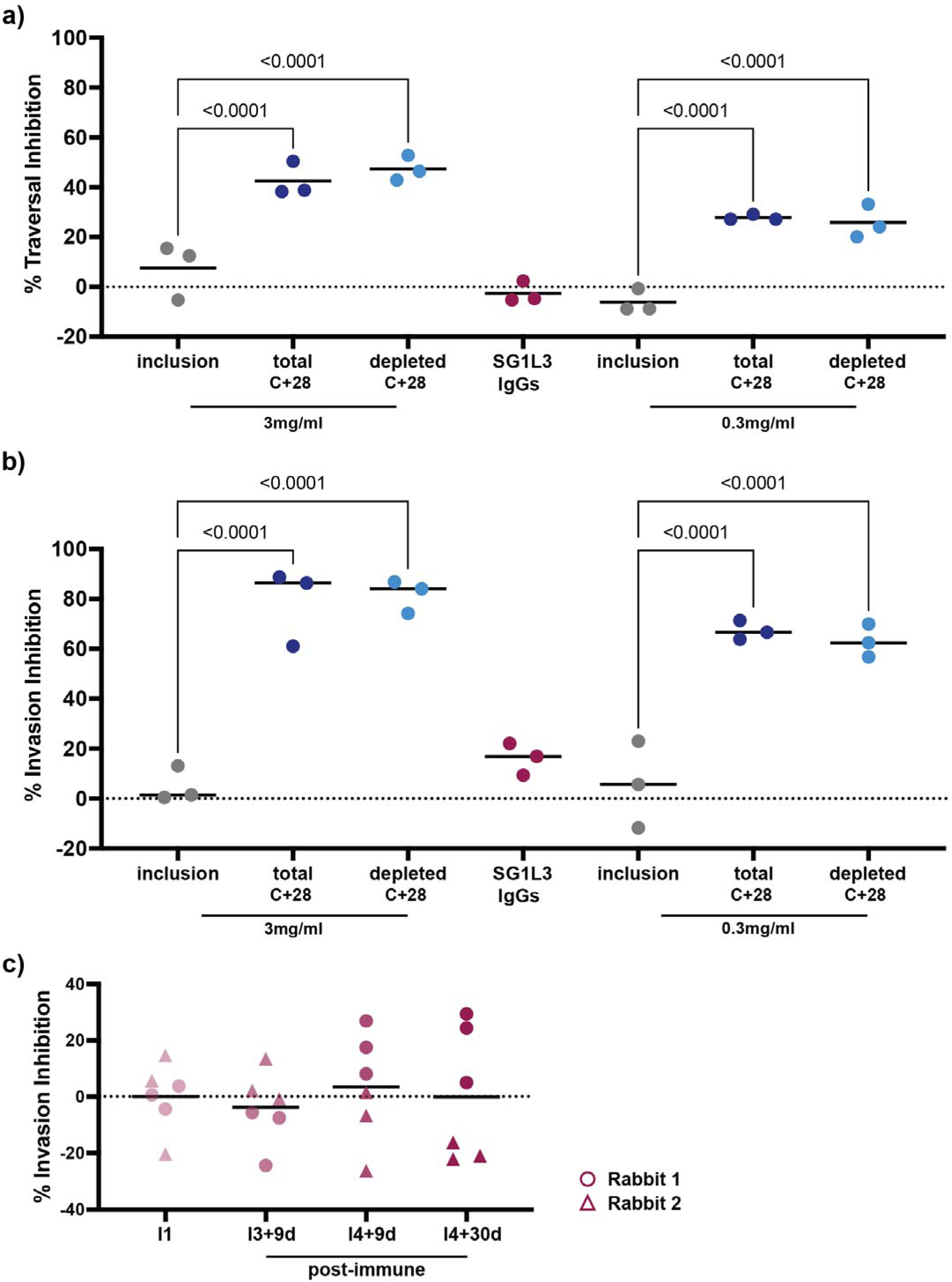
Polyclonal SG1L3 show no traversal or invasion inhibition of *P. falciparum* sporozoites **a)** Traversal inhibition and **b)** Invasion inhibition of HC-04 cells by *P. falciparum* sporozoites when incubated with inclusion (pre-immune), C+28 total and C+28 SG1L3 depleted (post-immune), at either 3mg/mL or 0.3 mg/mL IgGs from the immunized volunteer. Purified SG1L3-specific IgGs were reconstituted in a volume that matched the volume of the samples that were tested at 3 mg/mL. One-way ANOVA, with Šídák’s multiple comparisons test to its inclusion control. p>0.05 values are not shown. Invasion and traversal inhibition are calculated as the reduction in invaded/traversed HC-04 cells normalised to a no IgG control. **c)** Invasion inhibition of Pf sporozoites when incubated with purified polyclonal IgGs (2mg/ml) from SG1L3-immunized rabbits at different time-points. Bars represent mean.

We then proceeded with *in vitro* functional assays, and performed both traversal and invasion assays where *P. falciparum* sporozoites were incubated with total pre-immune IgG (inclusion), total IgG from C+28, SG1L3-depleted IgGs from C+28 and SG1L3-specific IgGs prior to invasion or traversal (Fig. 4a-b). We observed that both total and depleted IgGs, when tested at 3 mg/mL, were able to inhibit traversal and invasion of hepatocytes, whereas SG1L3-specific IgGs were not. Since the observed inhibition activity is very high and may be saturated, we further tested diluted antibodies at 0.3 mg/mL to test activity within the dynamic range. While the inhibitory activity dropped, SG1L3-depleted IgGs retained a similar inhibition of both traversal and invasion as total IgGs, showing that SG1L3-specific IgGs do not contribute to the non-CSP-mediated *in vitro* invasion and traversal activity in this volunteer.

We hypothesized that whole sporozoite immunization in humans may have elicited SG1L3 antibody levels too low to detect traversal or invasion inhibition and therefore immunized rabbits with recombinant SG1L3 to generate high SG1L3-specific antibody titres (Sup. Fig. 5a). Immunized rabbits successfully developed antibodies against recombinant SG1L3, able to recognize native SG1L3 in infected mosquito salivary gland lysate, as shown by ELISA and western blot (Sup. Fig. 5b-c). Post immunization IgGs were also able to recognize *P. falciparum* sporozoites, as detected by flow cytometry (Sup. Fig. 5d). Nonetheless, when purified IgGs from post immunization timepoints were incubated with *P. falciparum* sporozoites in an HC-04 invasion assay, they showed no inhibition (Fig. 4c).

**Figure 5:**
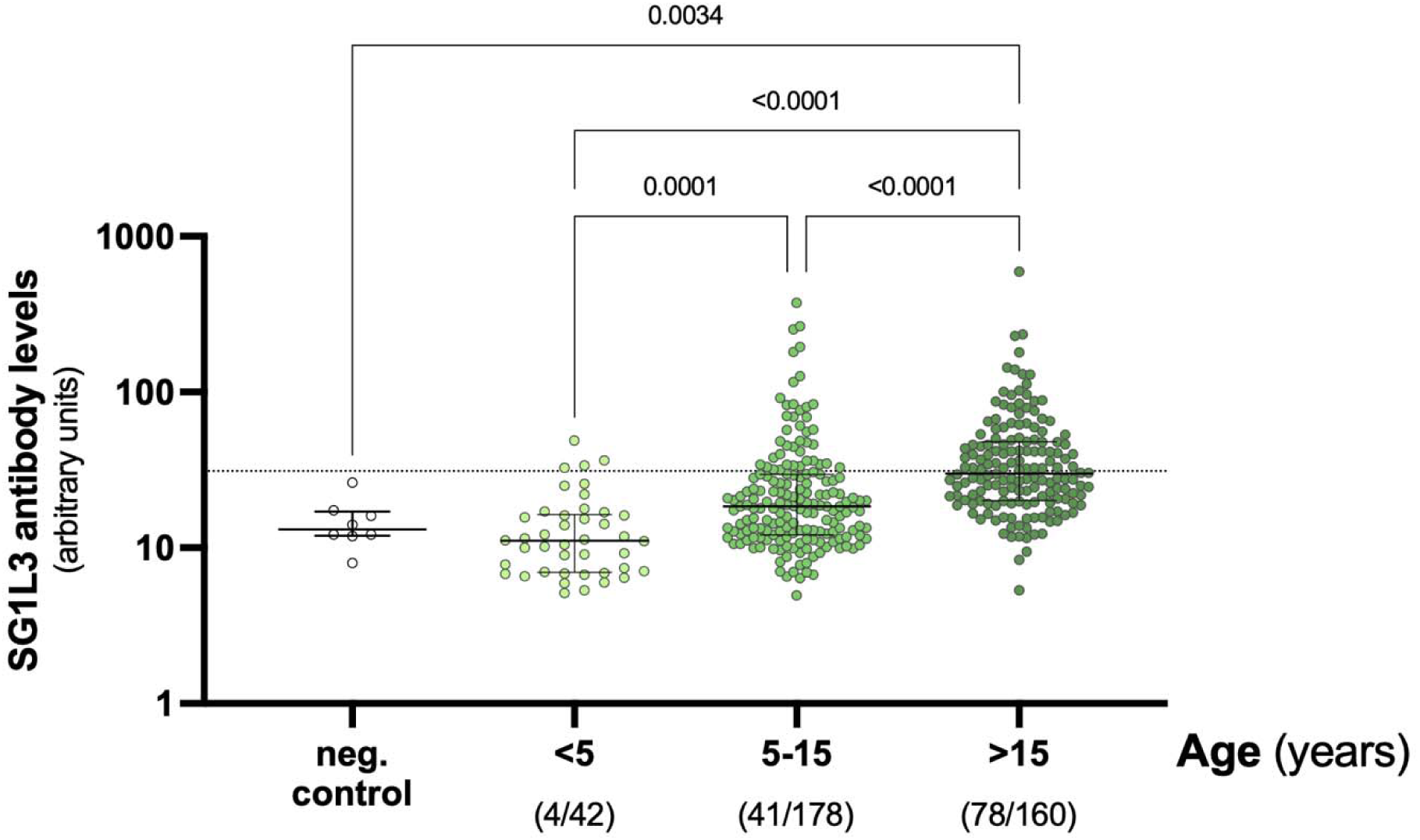
SG1L3 antibodies are present and increase with age in naturally exposed individuals from Burkina Faso. Recombinant SG1L3 salivary protein ELISA with plasma from Dutch malaria naïve volunteers (neg. control) or naturally exposed individuals from Burkina Faso, stratified by age: <5 years (n=42), 5-15 years (n=178) and >15 years (n=160). Dunn’s Kruskal-Wallis test with multiple comparisons. Bars indicate median ± IQR (interquartile range). p>0.05 values are not shown. Dashed line indicates positivity cut-off, defined as the average arbitrary units from the negative controls + 3xSD. The number of positive samples, and the number of total samples per age category, are shown below each group between brackets.

Taken together, our results show that polyclonal IgGs against salivary protein SG1L3 do not inhibit *P. falciparum* sporozoite invasion and traversal *in vitro*.

### Natural exposure to mosquito bites induces antibody responses against SG1L3

Considering that an experimental immunization by mosquito bites induced a strong antibody response to the salivary protein SG1L3, we wondered if antibody responses were detected under natural conditions among individuals regularly exposed to the bites of *Anopheles* mosquitoes. To this end, we recombinantly produced the orthologous *Anopheles gambiae* SG1L3 salivary protein and performed ELISAs to detect IgG responses in a sample set from individuals from an area of intense malaria transmission in Sapone health district, Burkina Faso (Collins, Ouedraogo et al. 2019, Collins, Ouedraogo et al. 2024). Exposure to *Anopheles* mosquitoes in the study area is high and primarily related to *Anopheles s.l.* (Guelbeogo, Goncalves et al. 2018). Antibody prevalence was 9.5% (4/42) in children <5 years of age, 23.0% (41/178) among children 5-15 years and 48.8% (78/160) among individuals older than 15 years. We further observed that the density of antibodies against salivary protein SG1L3 increased with age, possibly reflecting cumulative exposure to the bites of *Anopheles* mosquitoes (Fig. 5). Of note, we observed no significant difference in antibody levels between children under the age of 5, and Dutch blood bank controls, which suggests that antibody responses to the SG1L3 salivary protein may only become detectable after repeated or prolonged exposure to *Anopheles* bites. Our results show that salivary protein SG1L3 is able to induce antibody responses in humans after natural exposure to the bites of *Anopheles* mosquitoes.

## Discussion

Here, we present the first identification of non-CSP monoclonal antibodies (mAbs) from a participant who was successfully immunized under chemoprophylaxis. Previous work has characterized the polyclonal antibody repertoire elicited by natural infections and experimental immunizations with malaria sporozoites and identified antibody responses that target non-CSP antigens (Doolan, Southwood et al. 2003, John, Moormann et al. 2005, Doolan, Mu et al. 2008, Felgner, Roestenberg et al. 2013, Nahrendorf, Scholzen et al. 2014, Obiero, Campo et al. 2019) but never identified non-CSP mAbs. Using a newly developed target agnostic B-cell screening approach, we identified two mAbs that recognize *Anopheles stephensi* salivary protein SG1L3, and demonstrated that SG1L3 binds to the surface of sporozoites.

SG1L3 is not only the target of the isolated mAbs, but also one of the main antigens recognized by polyclonal antibodies from the same Dutch volunteer with strong functional invasion-inhibition in the absence of CSP antibodies (Fabra-Garcia, Yang et al. 2022). To assess whether SG1L3-specific antibodies from this volunteer indeed contributed to the invasion-inhibition activity, we tested the isolated mAbs and purified SG1L3-specifc polyclonal antibodies from this donor in an invasion inhibition assay and observed no activity. Importantly, we showed that depletion of SG1L3-specific antibodies did not affect the level of sporozoite traversal nor invasion inhibition, demonstrating that SG1L3 antibodies do not contribute to the observed invasion inhibition by CSP-depleted antibodies from this volunteer (Fabra-Garcia, Yang et al. 2022). As whole sporozoite immunization may have generated SG1L3 antibody levels too low to detect functional activity, we then tested if recombinant SG1L3 could generate functional antibodies in a rabbit immunization study and observed no inhibition of infection *in vitro.* We cannot rule out that our recombinant SG1L3 failed to induce functional antibodies due to glycosylation which could result in masking of protective epitopes, alter antigen processing and presentation, and promote immunodominant but non-neutralizing antibody responses (Rudd, Elliott et al. 2001) However, since none of our experiments with mAbs and polyclonal antibodies demonstrated functional activity SG1L3 antibodies, it seems unlikely that salivary protein SG1L3 is a target for liver stage infection.

Monoclonal antibody responses against *P. falciparum* CSP have been well described, where several studies have used recombinant PfCSP for B-cell sorting to identify potent monoclonal antibodies. These studies have used either B cells from naturally exposed individuals (Triller, Scally et al. 2017) or from immunized volunteers with either live attenuated sporozoites (Kisalu, Idris et al. 2018, Murugan, Buchauer et al. 2018, Tan, Sack et al. 2018, Wang, Pereira et al. 2020) or RTS,S vaccine (Oyen, Torres et al. 2017, Williams, Guerrero et al. 2024). Prior to this work, only two other studies used an antigen-agnostic method to search for targets of protection against the sporozoite stage of infection. The first study used immortalized memory B cells from Tanzanian volunteers immunised with live attenuated sporozoites (PfSPZ), and found only antibodies against CSP, highlighting CSP’s dominance in the immune response against *Plasmodium* (Tan, Sack et al. 2018). The second study target-agnostically isolated potent mAbs from naturally exposed and experimentally immunized volunteers that had polyclonal antibodies that recognized sporozoites but not recombinant PfCSP. However, the authors discovered that the novel mAbs they isolated recognized a cryptic epitope in CSP that is present on sporozoites, but not present in the recombinant PfCSP (Dacon, Moskovitz et al. 2025). While our study is the first study to successfully identify non-CSP mAbs from human volunteers as well as the target of these mAbs, we were not able to identify the target of the functional non-CSP antibodies described in our previous work from the volunteer studied here (Fabra-Garcia, Yang et al. 2022). We can rule out that the observed activity for non-CSP antibodies in this volunteer was mediated by antibodies targeting the cryptic epitope on CSP, as the CSP-depleted antibodies did not recognize native CSP in sporozoite lysate containing the cryptic epitope (Fabra-Garcia, Yang et al. 2022).

There are several other approaches that may enable the identification of the target(s) of the potent non-CSP antibodies from human volunteers. First, potential targets may be identified by profiling antibody responses after parasite exposure using a customized phage display such as PhIP-seq which contains peptides from both *P. falciparum* and *Anopheles* mosquitoes’ proteins (Raghavan, Kalantar et al. 2023), or protein microarrays (Felgner, Roestenberg et al. 2013, Obiero, Campo et al. 2019). Secondly, functional mAbs may be identified by leveraging single-cell RNA sequencing with paired B cell receptor (BCR) V(D)J profiling technology (Zheng, Terry et al. 2017). This technology could be used to compare B cell repertoires before and after controlled human infection with *Plasmodium falciparum*-infected mosquito bites, to enable identification of clonally expanded B cell populations. Thirdly, non-CSP-specific B cells could be isolated utilizing fluorescently labeled *Plasmodium falciparum* sporozoites, an approach that has been described against viruses (Doucett, Gerhard et al. 2005, Woda and Mathew 2015) and more recently for the identification of human mAbs against bacteria (van der Lans, Bardoel et al. 2024). While these methodologies may face challenges such as the identification of false positive targets (PhIP-seq/Microarray) or screening many irrelevant mAbs (single RNA sequencing) or being technically challenging, they may be attractive approaches that circumvent screening large numbers of single B cells and may therefore be more successful in identifying functional non-CSP targets.

Salivary proteins are essential for successful blood feeding by the mosquito vector. They play an important role in inhibiting coagulation and inflammation, and inducing vasodilation, among other roles (Barillas-Mury, Ribeiro et al. 2022). Mosquitoes inject saliva into the vertebrate host upon taking a blood meal, and recent studies have demonstrated that mosquito salivary proteins can either inhibit or enhance *Plasmodium* parasite infection (Arora, Chuang et al. 2023). Salivary proteins can play a role in the establishment of *P. falciparum* infection in the host in different ways. Salivary protein AgTRIO has been shown to enhance infection of murine *P. berghei* by influencing the local inflammatory response to sporozoites in the skin. However, when experiments are done *in vitro,* no influence in infection is observed, highlighting AgTRIO’s role in mediating the inflammatory response at the skin stage *in vivo* (Dragovic, Agunbiade et al. 2018, Chuang, Freudzon et al. 2019). On the other hand, salivary protein mosGILT has been shown to bind to the sporozoite surface and inhibit sporozoite gliding motility and cell traversal of *P. berghei* sporozoites *in vivo and in vitro*, suggesting that mosGILT reduces sporozoite infectivity (Schleicher, Yang et al. 2018). In the current study, we only assessed whether antibodies against SG1L3 could affect liver cell invasion and traversal, as the antibodies were obtained from a volunteer with strong liver cell invasion inhibition activity. Interestingly, in the study where mosGILT was found to bind to sporozoites and play a role in *P. falciparum* infection, SG1L3 salivary protein was also identified as one of many saliva proteins that were identified in mass spectrometry experiments with washed *P. falciparum* sporozoites (Schleicher, Yang et al. 2018). Using mAbs and recombinant SG1L3, we conclusively show in our study that SG1L3 binds to sporozoites. Since SG1L3 binds to sporozoites, it is tempting to speculate that SG1L3 may affect sporozoite infectivity during the skin stage of infection. Future studies should therefore assess the impact of (antibodies against) SG1L3 on sporozoite infectivity in skin models (Amino, Thiberge et al. 2006, Schleicher, Yang et al. 2018), or in mouse infection models (Dragovic, Agunbiade et al. 2018, Schleicher, Yang et al. 2018) that use mosquitoes to administer sporozoites to mice that are immunized with salivary protein SG1L3.

We found that SG1L3 salivary protein localizes to the median lobe of the mosquito salivary glands, similarly to other salivary proteins belonging to the SG1 family such as AgTRIO (Klug, Arnold et al. 2022) and saglin (O’Brochta, Alford et al. 2019, Klug, Arnold et al. 2022). Although sporozoites are known to preferentially infect the distal lateral lobes of the mosquito salivary glands (Wells and Andrew 2019), it is possible that saliva proteins and sporozoites interact when they come together in the salivary duct when mosquito probes for blood, possibly influencing subsequent infection outcomes.

Our work has shown that experimental immunization with three times 15 mosquito bites induces strong antibody responses against *A. stephensi* salivary protein SG1L3. Antibody responses against mosquito salivary proteins are an important avenue of research, since these are able to capture individual mosquito-human interactions, aiding the tracking of mosquito exposure in populations at risk of mosquito-borne diseases and evaluating vector control strategies success (Kearney, Heng-Chin et al. 2025). There are currently two mosquito salivary proteins, cE5 and SG6, that have shown promise as serological markers of exposure but are suboptimal. cE5 has shown to induce long-lived IgG responses, which limits utility for detecting short-term changes in mosquito exposure (Rizzo, Lombardo et al. 2014). SG6 has shown to induce immune tolerance, where individuals show no increase, or even a decline in antibody levels with increasing exposure to mosquito bites, which limits utility for populations in endemic areas (Rizzo, Ronca et al. 2011). Our results show that unlike SG6, salivary protein SG1L3 does not induce immune tolerance, since we observe an increase in antibody levels with cumulative exposure in increasing ages. Future studies should assess how long-lived SG1L3-induced antibodies are by following the individual antibody dynamics over time while monitoring mosquito exposure. This will also allow to ascertain if SG1L3 antibodies can be used as serological marker during low vector density, *e.g.* to distinguish mosquito exposure in different seasons or after introduction of vector control measures.

In this study, we used an agnostic single B-cell sorting approach that has allowed to expand the current knowledge of proteins present at the surface of *P. falciparum* sporozoites. By leveraging this knowledge, we identified a possible new marker to track mosquito exposure in malaria endemic populations and that may aid in elimination campaigns. Our study highlights the value of systematically characterizing the full humoral immune repertoire, including B cell responses to both parasite- and vector-derived antigens, to expand the pool of potential antibody targets and markers of mosquito exposure.

## Materials & Methods

### Study population and ethical approval

For characterization of non-CSP immune responses, PMBCs and plasma samples were obtained from Dutch malaria-naïve volunteers who participated in a chemoprophylaxis sporozoite (CPS) immunization trial (NCT02098590), approved by the Central Committee for Research Involving Human Subjects of The Netherlands (CCMO NL48732.091.14). Study protocol has been previously described (Walk, Reuling et al. 2017). Immunization scheme is shown in Supplementary Figure 1.

To describe naturally acquired antibody responses against salivary protein SG1L3, plasma samples were obtained from a previously described cohort study (Collins, Ouedraogo et al. 2019, Collins, Ouedraogo et al. 2024) conducted in Sapone, Burkina Faso, an area with intense and seasonal malaria transmission, in November 2019. The time of sampling coincided with the end of the transmission season. Study was approved by London School of Hygiene and Tropical Medicine ethics committee (14724), The Centre National de Recherche et de Formation sur le Paludisme institutional review board (2018/000002/MS/SG/CNRFP/CIB), and Burkina Faso national ethics committee for health research (2018-01-010).

### B-cell sorting and stimulation

PBMCs were rapidly thawed at 37°C, washed with 20% heat-inactivated FBS in RPMI with 0.0375 units/µl of Benzonase (Novagen, cat. no. 70664), and resuspended in 10% FBS in RPMI. Cells were counted with a haemocytometer using trypan blue exclusion dye, and allowed to rest for 1 hour at 37°C and 5% CO_2_. Cells were resuspended in 2% FBS in PBS, and stained for 1 hour at 4°C with CD19-BV421 (clone HIB19, BioLegend, cat. no. 30233), IgA PE (clone IS11-8E10, Miltenyi Biotec, cat. no. 130-114-002), IgM-APC/Cy7 (clone MHM-88, BioLegend, cat. no. 314520) and IgD-FITC (clone IA6-2, BioLegend, cat. no. 348206), washed and resuspended in 2%FBS in PBS. Cells were sorted using BD FACS Aria cell sorter, with 100 µm nozzle, following the gating strategy in Sup. Fig. 1c. Single B cells (1 cell/well) were sorted into 384 well-plates (Nunc MaxiSorp, cat. no. 242765) with Immunocult-XF T cell Expansion Medium (#10981, Stem Cell Technologies) supplemented with ImmunoCult™-ACF Human B Cell Expansion Supplement (#10974, Stem Cell Technologies). After sorting, B cells were spun (3min, 350xg) and incubated for 14 days at 37°C and 5% CO_2_. After 14-day stimulation, supernatants were collected into a new 384-well plate (Greiner, cat. no. 781201) with the help of a pipetting robot (INTEGRA Biosciences Assist Plus). B cells were immediately lysed in ice-cold lysis buffer (1M Tris-HCl (pH 8.0), RNase Inhibitor (New England Biolabs cat. no. M0314), in DEPC-treated water) according to previously published protocols (Huang, Doria-Rose et al. 2013), and stored at −80°C until further processing. Supernatants were immediately frozen at −80°C for later analyses in sporozoite lysate ELISA.

### B-Cell Receptor Amplification

From selected wells, we performed reverse transcriptase polymerase chain reaction (RT-PCR) following a previously established protocol (Tiller, Meffre et al. 2008). Briefly, cDNA was synthetized in nuclease-free water (Invitrogen cat. no. AM9920) with 1 µl 0.1M DTT (ThermoFisher Scientific), 0.5 µl dNTPs (10mM each, ThermoFisher Scientific cat. no. R0192), 1µl 50uM Random Hexamers (ThermoFisher Scientific), 1 µl 7% Igepal Ca-630 (Sigma Aldrich), 0.25 µl (8.5U) RNAsin inhibitor (Promega, cat. no. N2111), 0.25 µl (50U) Superscript III and 2.8 µl first strand buffer (Invitrogen, cat. no. 18080093). RT reaction was performed as follows: 42°C for 10 min, 25°C for 10 min, 50°C for 60 min and 94°C for 5 min. cDNA was stored at −20°C until further use. Using 3.8 µl of cDNA as a template, V gene transcripts were amplified separately for IgH, Igk and Igl following previously established protocols (Gieselmann, Kreer et al. 2021). Briefly, a master mix was prepared, that contained 0.09µl Platinum Taq DNA Polymerase, 2.05 µl Platinum Taq PCR buffer, 1.23 µl KB extender, 0.61µl MgCl_2_ (all Invitrogen, cat. no. 10966018), 0.41 µl 10 mM dNTP mix (Thermo Scientific, cat no. R0192), 0.09 µl forward and 0.09 µl reverse primers (50 µM), and 14.68 µl nuclease-free water DEPC treated water (Invitrogen cat no. AM9920). Reaction was run as follows: 94°C 2min, 94°C 30s, 57°C 30s,72°C 55s, for 50 cycles, and a final step at 72°C 6min. Second PCR was identical to the first PCR, using second PCR-specific primers, and using 3µl of unpurified first PCR product as template. Second PCR reaction is as follows: 94°C 2min, 94°C 30s, 57°C 30s, 72°C 45s, for 50 cycles, and a final step at 72°C 6min. PCR products were analysed by 1% agarose gel electrophoresis in 1x TBE buffer, to evaluate which chains were successfully amplified. PCR products for heavy and light chains were expected at 500 bp or 450 bp, respectively. Primers used in this work have been previously described (Gieselmann, Kreer et al. 2021).

PCR products with expected size were sent for Sanger Sequencing (BaseClear, the Netherlands), and sequencing results were uploaded to IGMT V-Quest (Brochet, Lefranc et al. 2008, Giudicelli, Brochet et al. 2011) to annotate heavy- and light-chain sequences. Sequences that encoded productive variable domains were selected for cloning into expression vectors.

### BCR cloning

For cloning variable domains into expression vectors, a 3rd PCR was performed on products from the first PCR, following previously described protocol (Gieselmann, Kreer et al. 2021). Briefly, a master mix was prepared, that contained 0.5µl Q5 high-fidelity DNA polymerase (New England Biolabs, cat. no. M0491S), 10µl Q5 PCR buffer, 10µl Q5 high GC enhancer, 1µl dNTP mix and 24.1µl nuclease-free water. 0.5µl forward primer and 0.5µl reverse primer was added, specific for heavy, kappa or lambda chain. 1µl of unpurified first PCR product was used as a template and third PCR reaction was run as follows: 98°C, 30s, 98°C 10s, 72°C 45s, for 35 cycles, 72°C 2min. PCR products were analysed on a 1% agarose gel in 1x TBE buffer. Positive chains were purified using the QIAquick PCR purification kit (Qiagen, cat. no. 28104), and concentrations were adjusted to 40ng/µl with nuclease-free water. IgH, Igk and Igl expression vectors (Tiller, Meffre et al. 2008) were linearized using the following restriction enzymes, respectively: EcoRI-HF and SalI-HF, EcoRI-HF and BsiWI-HF, and EcoRI-HF and XhoI (all restriction enzymes from New England Biolabs). Reactions were set up as follows: 10µg vector was incubated for 16h at 37°C with 2.5µl per enzyme (50U), 5µl CutSmart Buffer (10x) and completed to 50µl with nuclease-free water. Vectors were purified using the QIAquick PCR purification kit and concentrations were adjusted to 80ng/µl with nuclease-free water.

For cloning individual variable regions into the expression vectors, the following reaction was set up on ice: 0.2µl T4 DNA polymerase (New England Biolabs, cat. no. M0203S), 1µl NEBuffer 2.1, 1µl linearized expression vector and 6.8µl nuclease-free water. 1µl of purified heavy, kappa or lambda chain was added and reactions were incubated at 24°C for 2.5min, followed by a 10min incubation on ice. For transformation, 4µl was transferred to 40µl chemically competent *E. coli* Stellar cells and incubated on ice for 30 min. The reaction was incubated at 42°C for 45s (heat shock), followed by incubation on ice for 2min. 50µl of Lysogeny Broth (LB) medium was added and reactions were incubated at 37°C, 210 rpm for 1h. Reactions were briefly centrifuged, streaked with glass beads on LB-ampicillin agar plates and plates were incubated at 37°C for 16 h.

Colonies were examined by colony PCR. Colonies were inoculated in 5µl Milli-Q and incubated at 95°C for 10min. A master mixed was prepared that contained 0.125µl DreamTaq DNA Polymerase (ThermoFisher Scientific, cat. no. EP0712), 2.5µl DreamTaq Green Buffer, 0.2µl dNTP mix and 16.175 µl nuclease-free water. 0.5 µl forward primer and 0.5 µl reverse primer were added and PCR mix was added to colonies. PCR reaction was run as follows: 95 °C, 3 min, 95 °C 30 s, 54 °C 30 s, 72 °C 1 min, for 35 cycles, and a final step at 72 °C 5 min. PCR products were analysed on a 1% agarose gel in 1x TBE buffer, and positive colonies were grown in 5 ml LB broth medium containing ampicillin at 37 °C and 210 rpm for 16 hours.

Plasmids were isolated using the Qiaprep Spin Miniprep Kit (Qiagen, cat. no. 27104) and sent for Sanger Sequencing (Baseclear, The Netherlands). Sequencing results were uploaded to IGMT V-Quest to check if chains were productive, and aligned to second PCR product to confirm no mutations were introduced. Primers used here have been previously described by (Gieselmann, Kreer et al. 2021).

### Recombinant antibody expression

Antibodies were expressed in HEK293T cells. HEK293T cells were cultured in DMEM, high glucose (ThermoFisher Scientific, cat. no. 41965062) supplemented with 10% heat-inactivated FBS and 20U/ml penicillin/streptomycin (ThermoFisher Scientific, cat. no. 15140122) at 37°C and 5%CO_2_. One day before transfection, 4×10^6^ cells were seeded per T75 flask. 3 hours prior to transfection, cells were washed with 1x PBS and 10 mL fresh medium was added in which FBS was replaced with 10% ultra-low IgG FBS (ThermoFisher Scientific, cat. no. A3381901). Transfection mix was prepared as follows: DNA mix was added at 1 µg per mL culture, with a heavy chain:light chain ratio of 2:1. Heavy and light chains were mixed in Opti-MEM (ThermoFisher Scientific, cat. no. 11058021). Polyethylenimine (PEI) (Polysciences, cat. no. 23966) was diluted in Opti-MEM, with DNA:PEI ratio of 1:3. DNA mix was added to the PEI mix drop-by-drop, and mixes were incubated for 30 min in the dark. 1ml transfection mix was added to the T75 flask drop-by-drop. Flasks were gently shaken and incubated at 37°C and 5% CO_2_ for 6-7 days.

### ELISA

Sporozoite lysate was prepared as followed: Sporozoites were isolated from mosquitoes’ salivary glands, pelleted and stored at −80°C. For lysis, 1×10^6^ sporozoite pellet was resuspended in 100ml lysis buffer (150 mM NaCl, 20 mM Tris-HCl, 1% triton, 1 mM EDTA, pH 7.5) and 1 µl protease inhibitor (ThermoScientific cat. no. 78410), incubated on ice for 15 minutes. Supernatant was then collected after 10min spin at max speed in a tabletop centrifuge, and stored at −80 °C until further use.

Recombinant proteins (1-2 µg/mL) or *P. falciparum* sporozoite lysate (3125 spz/well) were coated in 1x PBS overnight at 4°C, in 96-well plates (Nunc MaxiSorp cat. no. 439454). Plates were then washed with 1x PBS, and blocked (5% milk in 1x PBS) for 1 hour at room temperature (RT). Supernatants from stimulated B cells (1:5 dilution), volunteer plasma (1:200 dilution) or monoclonal antibodies were then added to the plates, and incubated for 3 hours at RT. Plates were then washed thrice with 0.05% Tween 20 in 1x PBS, and incubated with anti-human IgG-HRP (Pierce cat. no. 31412) for 1hour at RT. Plates were then washed thrice with 0.05% Tween 20 in 1x PBS, and thrice in 1x PBS, and incubated with TMB (Surmodics BioFX TMB, cat. no. TMBW-1000-01) for 10-15 minutes. Reaction was stopped with the addition of 0.2M H_2_SO_4_ and read on iMark Microplate Reader (Bio-Rad) at 450 nm optical density.

For the determination of antibody responses against SG1L3, analyses of ELISA results were performed using Auditable Data Analysis and Management System for ELISA (ADAMSEL FPL v1.1). For determination of arbitrary units, a serial diluted pool of serum from 15 individuals who had high reactivity against SG1L3 was used. In order to convert optical density (OD) units into arbitrary units (AU), undiluted control serum was defined to contain 100 AU.

### Immunoprecipitation

Target of mAbs was identified by immunoprecipitation using Dynabeads^TM^ Protein G Immunoprecipitation Kit (Invitrogen, cat. no. 10007D), following manufacterer’s instructions, with the following adaptations: 10 µg of 16O7 mAb was coupled to DynaBeads^TM^, and crosslinked to beads using the BS3 (bis(sulfosuccinimidyl)suberate) buffer (Invitrogen, cat. no. A39266). Beads with coupled 16O7 were incubated with lysate of 46 uninfected *Anopheles stephensi* mosquito salivary glands, lysed in 1% Triton and 1x protease inhibitor (ThermoScientific cat. no. 78410). Proteins binding to 16O7 beads were eluted in 5% SDS in PBS, followed by 10 minutes incubation at 70°C.

### Mass spectrometry analysis

Samples were processed on S-TRAP micro (Protifi). Samples were adjusted to 5% SDS final concentration. Reduction and alkylation were performed with 10 mM TCEP and 50 mM iodoacetamide. Samples were then acidified with phosphoric acid (stock 12%) to 1.2% final concentration and then diluted (1:7 ratio) with 90% methanol/100 mM TEAB. Samples were then transferred onto the S-TRAP spin columns and spun through with 4000 xg and washed 4 times with 150 µl 90% methanol/100mM TEAB (4000 xg). 1.5 µg trypsin (Promega) in 20 µl 50mM TEAB was then added to the spin column and was digested over night at 37 °C. No shaking was applied. Next day, peptides were eluted with 150 µl 0.1% formic acid and 150 µl 50% acetonitrile/ 0.1% trifluoroacid and then dried to total dryness in centrifugal evaporator.

Peptides were resuspended in 5% formic acid and 5% DMSO and then trapped on an Acclaim™ PepMap™ 100 C18 HPLC Columns (PepMapC18; 300 µm x 5 mm, 5 µm particle size, Thermo Fisher) using solvent A (0.1% Formic Acid in water) at a pressure of 60 bar and separated on an Ultimate 3000 UHPLC system (Thermo Fisher Scientific) coupled to a QExactive mass spectrometer (Thermo Fisher Scientific). The peptides were separated on an Easy Spray PepMap RSLC column (75µm i.d. x 2µm x 50mm, 100Å, Thermo Fisher) and then electro-sprayed directly into an QExactive mass spectrometer (Thermo Fisher Scientific) through an EASY-Spray nano-electrospray ion source (Thermo Fisher Scientific) using a linear gradient (length: 60 minutes, 5% to 35% solvent B (0.1% formic acid in acetonitrile and 5% dimethyl sulfoxide), flow rate: 250 nL/min). The raw data was acquired in the mass spectrometer in a data-independent mode (DIA). Full scan MS spectra were acquired in the Orbitrap (inclusion list with scan range 495 to 995 m/z, 20m/z increments, with an overlap of +/- 2 Daltons, resolution 35000, AGC target 3e6, maximum injection time 55ms). After the MS scans peaks were selected for HCD fragmentation at 28% of normalised collision energy (nce). HCD spectra were also acquired in the Orbitrap (resolution 17500, AGC target 1e6, isolation window 20m/z).

The acquired raw MS data were searched in a library-free search in DIA-NN [version1.8.1] *Anopheles stephensi* protein database using the database *A. stephensi* SDA-500 release 59 from vectorbase.org. FDR (false discovery rate) was set to 0.01 and MBR (match between runs) was enabled. Cross run normalisation was switched on. Cross run normalization was performed by MaxLFQ, a widely used generic method for label-free quantification. Following modifications were applied, carbamidomethyl (C) and oxidation (M).

### Western Blot

Sporozoites were collected from hand-dissected salivary glands of *P. falciparum* NF54 infected mosquitoes. Sporozoites were spun down (16,000 xg, 5 minutes) and resuspended in 1x LDS (GenScript, cat. no. M00676), denatured for 10 minutes at 70 °C. Proteins were separated by electrophoresis on a 4-12% Bis-Tris SurePAGE (GenScript, cat. no. M00654) or NuPAGE (Invitrogen, NP0326BOX) Polyacrylamide gel and precision plus dual color standard (Bio-Rad, cat. no. 1610374) was used as a protein marker. Proteins were then transferred to a nitrocellulose membrane using the Trans-Blot Turbo Transfer System (Bio-Rad). Membrane was washed once in 1x PBS, and then blocked with 5% milk PBS, for 1 hour at RT, or overnight at 4 °C. After blocking, membrane was incubated with different primary antibodies, either mAbs (2 µg/ml) or CaptureSelect™ anti-C-tag biotin conjugate (Thermo Scientific, cat. no. 7103252100, 1:1000) in 1% milk in 0.05% Tween-20 in PBS for 1 hour at RT. Following 3 washes with 0.05% Tween-20 in PBS, membranes were incubated with secondary antibody, anti-human IgG HRP (Pierce, cat. no. 31412, 1:10,000) or streptavidin HRP (R&D Systems, cat. no. 890803, 1:2000) diluted in 1% milk in 0.05% Tween-20 in PBS. After 3 washes with 0.05% Tween-20 in PBS, and once washed with 1x PBS, membranes were developed by adding Clarity Max ECL substrate (Bio-Rad). Images were acquired using ImageQuant LAS4000 (Bio-Rad). A protein marker WesternSure pen (LICORbio cat. no. 926-91000) was used to visualise the ladder.

### Recombinant protein expression

Codon-optimized constructs for expression of SG1 family proteins, containing a C-tag (EPEA), in S2 cells or HEK293T were ordered from GeneArt Gene Synthesis service (Thermo Fisher Scientific).

For expression in S2 cells, inserts were sub-cloned into a modified pExpreS_2_-2 plasmid (ExpreS^2^ion Biotechnologies, Denmark) to include an N-terminal Kozak sequence (GCCACC), BiP insect signal peptide (MKLCILLAVVAFVGLSLG) and an hexahistidine tag (HHHHHH). *Drosophila melanogaster* Schneider’s S2 cells (ExpreS2ion Biotechnologies, Denmark) were maintained in EX-CELL420 medium (Sigma-Aldrich) in shaker flasks at 25 °C. For transfection, 2×10^6^ cells/mL were transferred to T12.5 flask, and 25 μL of ExpreS2 Insect-TR 5x transfection reagent (ExpreS2ion Biotechnologies, Denmark) and 6.25 μg of plasmid DNA was added to the cells. After 2-4 hours, culture was supplemented with addition of 10% FBS. The next day, geneticin (G418) was added (4 mg/mL) for selection of successfully transfected cells. Cells were sub-cultured every 3-4 days, and maintained at a density of 1×10^6^ cells/mL. Antibiotic pressure was maintained for 24 days to remove all non-transfected cells. After 24 days, stably transfected cells were maintained in shake flaks at 115 rpm, in EX-CELL240 medium, sub-cultured every 4 days to 8×10^6^ cells/mL. Supernatant was harvested at each sub-culture, centrifuged at 3000 xg and stored at −20 °C until use.

SG1L3 protein expression in S2 cells failed and was transiently expressed in HEK293T cells instead. To this end, a codon-optimized construct encoding SG1L3 in a pcDNA3.4-TOPO vector (Thermo Fisher Scientific), with an N-terminal Kozak sequence (GCCACC), an hexahistidine tag (HHHHHH) and a c- tag (EPEA) was ordered. For expression of SG1L3 in HEK293T cells, the same protocol was used as for mAb expression, at 1:5 DNA:PEI ratio.

### Affinity purification and size exclusion chromatography

Supernatants containing proteins of interest were thawed and centrifuged at 3000 xg, and filtered using a 0.45 µm PES filter unit (Avantor). Supernatants were then concentrated using Vivaflow200 membrane (Sartorius). Samples were then loaded onto a 1mL CaptureSelect™ C-tag XL column (ThermoScientific, cat. no. 494307201) on the ÄKTA start (GE Healthcare). The column was then washed (20mM Tris in MilliQ, pH7.2-7.4) and the captured proteins were eluted (20mM Tris, 2.0M MgCl_2_ in MilliQ, pH7.2-7.4). Fractions that contained the target protein were pooled and dialysed overnight at 4°C using a 3.5 kD membrane (Spectrum) in 1x PBS. The next day, sample was concentrated using a 10 kD MWCO protein concentrator (ThermoScientific).

Samples were further purified by size exclusion chromatography (Amersham Biosciences). Samples were loaded on a Superdex200 Increase 10/300 GL size exclusion column (GE Healthcare), using 1x PBS (pH 7.2-7.4) as running buffer. Fractions containing purified protein were assessed by Coomassie stained SDS-PAGE gel, pooled and concentrated using a 10 kD MWCO protein concentrator (ThermoScientific). Concentration was measured on Nanodrop 1000, snap-frozen in liquid nitrogen, and stored at −80 °C until further use. SG1L3 protein showed signs of aggregation when using PBS, and therefore 0.02% Empigen^®^ BB (Sigma-Aldrich) was added to all buffers during purification. To analyse glycosylation in the recombinant proteins, Pierce™ Glycoprotein Staining kit (Thermo Fisher, cat. no. 24562) was used, following manufacturer’s instructions

### Immunofluorescence Microscopy

For *Anopheles stephensi* salivary gland immunofluorescence microscopy, a previously described method was used, with small modifications (Wells and Andrew 2015). Briefly, head with attached salivary glands from *Anopheles stephensi* mosquitoes were freshly hand-dissected in 1x PBS and stored in an Eppendorf tube on ice. Salivary glands were fixed in ice cold absolute acetone for 90 seconds at RT. Acetone was then removed, and 1x PBS was added, and salivary glands were incubated at RT for 30 minutes. 10 µg/ml primary antibody was added and incubated for 3 hours at RT. Salivary glands were washed thrice, with 5 minutes incubation between washes. Goat anti-human Alexa Fluor 488 secondary antibody (Invitrogen, cat. no. A11013) was added and incubated overnight at 4 °C. Salivary glands were then incubated with DAPI (Invitrogen, cat. no. D1306) for 30 minutes at RT, washed thrice with 1x PBS. Salivary glands were then freed from the mosquito head in a 100% glycerol solution in a microscopy slide, and a coverslip was mounted on top. Slides were kept at 4 °C until imaging.

For sporozoite immunofluorescence microscopy, 50,000 sporozoites were allowed to settle for 20 minutes on 12 mm poly-L-Lysine treated coverslips (Corning, cat. no. 354085) in a 24-well plate. Sporozoites were then fixed with 4% formaldehyde (Thermo Scientific cat. no. 28906) in 1x PBS for 10 minutes, washed once with 1x PBS and permeabilized with 0.1% triton in PBS for 10 minutes. Following three washes with 1x PBS, sporozoites were blocked in 3% BSA in PBS for 1 hour. Primary antibodies (10 µg/ml) were added diluted in 3%BSA in PBS, and incubated for 1 hour, followed by three washes in 1x PBS. Secondary antibodies goat anti-human IgG Alexa fluor 488 (Invitrogen, cat. no. A11013) and goat anti-mouse Alexa Fluor 568 (Invitrogen, cat. no. A11031) were added diluted in 3%BSA in PBS and incubated for 1 hour. Following three washes with 1x PBS, sporozoites were stained with 3 µM of DAPI (Invitrogen, cat. no. D1306) for 5 minutes, washed once with 1x PBS and once with water and coverslips were mounted onto microscopy slides using Fluoromount G mounting medium (Invitrogen, cat. no. 00-4958-02). All incubations were performed at room temperature. Images were acquired using LSM 900 confocal (Zeiss) using 40x oil (salivary glands) or 63 x oil (sporozoites) objectives. Images were processed using ImageJ (NIH) software.

### Sporozoite Binding assay by Flow Cytometry

Salivary gland *P. falciparum* NF54 sporozoites were pelleted and resuspended in 1× PBS, and incubated with 1 µM Syto61 Red Fluorescent Nucleic Acid Stain (Invitrogen) for 30 minutes at 4 °C. Following incubation, the sporozoites were pelleted at 16,000 g for 5 minutes, and resuspended in 1x PBS. For incubation with monoclonal antibodies, sporozoites were plated at 5×10^4^ sporozoites per well in a v-bottom plate, and incubated with the monoclonal antibodies (10 µg/mL in 3% w/v BSA/PBS) for 30 minutes at 4 °C. Sporozoites were then incubated with goat anti-human IgG Alexa Fluor 488 (Invitrogen cat. no. A11013, Thermo Fisher Scientific, 1:200) in 3% BSA/PBS for 30 minutes at 4°C. For incubation with recombinant proteins, sporozoites were then plated at 5×10^4^ sporozoites per well on a v-bottom plate (Costar), and incubated with the recombinant proteins (10 µg/mL in 3% w/v BSA/PBS) for 30 minutes at 4 °C. Sporozoites were then incubated with a 1:200 dilution of CaptureSelect™ anti-C-tag biotin conjugate (Thermo Scientific, cat. no. 7103252100) in 3% BSA/PBS for 30 minutes at 4 °C, and subsequently incubated with 1:200 Streptavidin Alexa Fluor 488 (Invitrogen, cat. no. S11223, Thermo Fisher Scientific) in 3% BSA/PBS for 30 minutes at 4 °C. For both assays, sporozoites were then fixed in 1% paraformaldehyde (Thermo Scientific, cat. no. 28906). Between protein and all antibody incubations, sporozoites were washed with 1x PBS, followed by centrifugation at 3220 × g for 3 minutes. Samples were acquired by flow cytometry using Gallios cytometer (Beckman Coulter), using Kaluza 1.0 software (Beckman Coulter). FlowJo v10.10.0 (BD Life Sciences) was used for data analyses.

### Gamete Binding Assay

Mature female gametes were obtained from heparin-treated stage V gametocyte cultures collected between 13 and 16 days after induction. Cells were concentrated by resuspension in fetal calf serum (FCS) at 50% of the initial culture volume, and gametogenesis was triggered by incubation at room temperature for 30 min under continuous agitation. Following induction, cells were pelleted by centrifugation at 2,000 × g at 4 °C, resuspended in PBS, and purified by density gradient centrifugation using a 12.4% Nycodenz cushion. Gradients were centrifuged at 3,500 × g for 30 min at 4 °C without deceleration, after which the fraction enriched for female gametes was recovered and washed in PBS. Purified gametes were distributed into non-treated, V-bottom 96-well plates (Costar) at a density of 5 × 10 cells per well. All incubation steps were performed at room temperature in PBS supplemented with 2% FCS (Gibco) and 0.02% sodium azide, with washes carried out using ice-cold PBS. To enable exclusion of non-viable cells, the fixable viability dye eFluor™ 780 (Invitrogen; 1:1000) was included in all secondary antibody staining steps. For analysis of binding by recombinant proteins, gametes were incubated with recombinant proteins for 1 h (10 µg/ml), followed by incubation with a mouse anti-His tag primary antibody (Sigma-Aldrich, cat. no. A7058; 1:200). Binding was detected using a chicken anti-mouse Alexa Fluor 488 (Invitrogen cat. no. A21200, Thermo Fisher Scientific, 1:200), for 30 min. In antibody-binding assays, gametes were incubated for 1 h with human antibodies at a final concentration of 1 μg/mL, washed, and labeled with a goat anti-human Alexa Fluor 488 secondary antibody (Invitrogen, A11013; 1:200). Following staining, samples were resuspended in PBS and analyzed by flow cytometry on a Gallios™ 10-color cytometer (Beckman Coulter), analyses were restricted to viable gametes.

### Traversal and Invasion Inhibition Assays

One day prior to assay, hepatocytes from HC-04 hepatoma cell line were seeded at 5×10^4^ cells/well in a flat-bottom 96-well plate (Corning, Merck), previously coated with 50 µg/mL rat-tail collagen (Sigma-Aldrich). Cells were grown in DMEM/ F12 supplemented with L-Glutamine, 5% FCS, 100 U/mL penicillin/streptomycin (all Gibco) at 37°C with 5% CO_2_. Salivary gland dissected NF54 *P. falciparum* sporozoites were pre-incubated on ice for 30 minutes with human malaria naive heat-inactivated serum, monoclonal (100 µg/ml) or polyclonal antibodies (2-3mg/ml), 50 µg/mL 2A10 or 1.25 µg/mL Cytochalasin D (Sigma Aldrich cat.no. C2618). For traversal, sporozoites were incubated with 1.25 mg/mL dextran prior to addition to the hepatocytes (Sigma-Aldrich, cat. no. FD10S). 4×10^4^ (traversal) or 5×10^4^ (invasion) sporozoites were added per well and incubated for 2.5-3 hours at 37°C and 5% CO_2_. After incubation, cells were washed thrice with 1x PBS, and detached using 0.25% Trypsin-EDTA (Gibco). Trypsin was inactivated by addition of 10% heat-inactivated human serum, and cells were transferred to a V-bottom plate, where they were washed again. For traversal assay, cells were immediately fixed in 1% paraformaldehyde (ThermoScientific) measured on Gallios flow cytometer (Beckman Coulter) the same day. For invasion assay, cells were stained with 1:2000 exclusion dye LifeDead780 eFluor (Invitrogen) for 20 minutes at 4°C, then permeabilized with eBioscience™ Foxp3 / Transcription Factor Fixation/Permeabilization (Invitrogen) for 30 minutes at 4°C. Cells were then stained intra-cellularly with 3SP2 antibody conjugated to Alexa Fluor 488 for 30 minutes at 4°C. Cells were fixed in 1% paraformaldehyde (ThermoScientific) and acquired on Gallios flow cytometer (Beckman Coulter) the next day, using Kaluza 1.0 software (Beckman Coulter). FlowJo software (V10.10.0, BD Life Sciences) was used for analysis.

### Polyclonal IgG purification

From the human plasma, IgGs were purified using saturated ammonium-sulphate precipitation. Plasma was diluted 1:1 in Mili-Q water, and ammonium-sulphate (Thermo Fisher Scientific, cat no. 45216) was added to 50% final concentration, and incubated for 15-30 minutes at RT. Sample was centrifuged at 16,100 x g at RT for 10 minutes. Pellet was resuspended in 50% ammonium sulphate and centrifuged at 16,100 x g at RT for 10 minutes. Pellet was dissolved in 1xPBS, spun down at 16,100 x g at RT for 10 minutes, and the supernatant filtered through a 0.45 µm filter. IgGs were then purified by 1mL Hi Trap protein G columns (GE Healthcare Life Science) on the ÄKTA start (GE Healthcare). 1X PBS and pH 2.8 amine-based IgG elution buffer (Pierce cat. no. 21004, Thermo Fisher Scientific) were used as binding and elution buffers, respectively. Eluted fractions were neutralized with 1M Tris pH 8.8 (Serva, cat. no. 3974.01). Purified IgGs were then buffer exchanged to 0.25x PBS using a centrifugal concentrator (Amicon Ultra-15, 30 kDa; Merck Millipore). Total IgG concentration was measured using NanoDrop 1000 (Thermo Fisher Scientific)

From the rabbit serum, 330 µL rabbit serum was diluted in 170 µL 1x PBS, and then bound to a Cytiva Ab SpinTrap column (Cytiva), incubated for 4 minutes at RT with occasional mixing. Sample was then centrifuged twice for 30 s at 100 g, RT, and column was washed with 1x PBS. IgGs were then eluted in a pH 2.8 amine-based IgG elution buffer (Pierce cat. no. 21004, Thermo Fisher Scientific). Eluted fractions were neutralized with 1M Tris pH 8.8 (Serva, cat. no. 3974.01a). Purified IgGs were then buffer exchanged to 0.25x PBS using 30K MWCO Pierce Concentrator (Pierce cat. no. 88502, Thermo Fisher Scientific).

### SG1L3-specific antibodies depletion and purification

In order to deplete and purify SG1L3-specific antibodies specific from the immunized volunteer, an affinity column was created by coupling 0.8 mg of recombinant SG1L3 to N-hydroxysuccinimide (NHS)-activated high performance affinity chromatography column (HiTrap™ NHS-activated HP 1 mL, GE Healthcare), according to manufacturer’s instructions.

For depletion and purification of SG1L3-specific antibodies, the SG1L3 NHS column was equilibrated with 1x PBS, before total IgG from volunteer plasma was added. The flow-through was collected, and the column washed with 1x PBS, followed by elution of SG1L3-specific IgGs with the addition of IgG elution buffer (Pierce cat. no. 21004, Thermo Fisher Scientific). The purification was repeated using the flow-through as the starting material. The final flow-through (SG1L3-depleted fraction) and both eluates (SG1L3-specifc IgG fraction) were buffer exchanged to 0.25x PBS and concentrated using a using a centrifugal concentrator (Amicon Ultra-15, 30 kDa; Merck Millipore). Depletion of SG1L3-specific antibodies was confirmed by ELISA and Western Blot (Supplementary Figure 3).

### Rabbit immunizations with recombinant SG1L3

Rabbit immunizations with recombinant SG1L3 were performed by Eurogentec (Kaneka Eurogentec S.A, Belgium) using the classical immunization program, following the immunization scheme illustrated in Supplementary Figure 4 a. Rabbits were immunized 4x with 100 µg recombinant SG1L3 per injection with complete Freund’s adjuvant (first vaccination) and incomplete Freund’s adjuvant (booster vaccinations).

### Statistical Analysis

All statistical analyses were performed using GraphPad Prism software version 10.6.1. The specific statistical tests used in each experiment are described in the corresponding figure legends. Statistical significance was defined as P value of ≤ 0.05.

## Acknowledgements

We thank the volunteers that participated in our study, both from the Netherlands and Burkina Faso. Mosquito breeding, infections and dissections were performed by Jolanda Klaassen, Laura Pelser-Posthumus, Astrid Pouwelsen, and Jacqueline Kuhnen. We acknowledge the support of the Radboudumc Technology Centers for Flow cytometry and Microscopy. Expression vectors for antibody cloning were kindly provided by Prof. Hedda Wardemann.

## Funding

This work was supported by the Bill and Melinda Gates Foundation (Grant INV-004976). C.M.A is further supported by a NWO Veni Fellowship (VI.VENI.022.381) and M.M.J. by a NWO Vidi Fellowship (VI.VIDI.192.061).

## Declaration of Interests

Authors declare no conflict of interests.

## Author Contributions

Conceptualization: C.M.A.; A.F.G.; R.W.S.; M.M.J.

Investigation: C.M.A.; R.C.v.D.; A.F.G; S.G.; G.-J.v.G.; K.T.; S.H.; R.S.; M.v.d.V.-B.

Formal analysis: C.M.A.; M.M.J. Visualization: C.M.A. Resources: C.D.; A.B.T.;

Writing—original draft: C.M.A.; M.M.J. Writing—review and editing: all authors Supervision: R.W.S.; T.B.; M.M.J. Funding acquisition: R.W.S.; M.M.J.

---

**Supplementary Figure 1:**
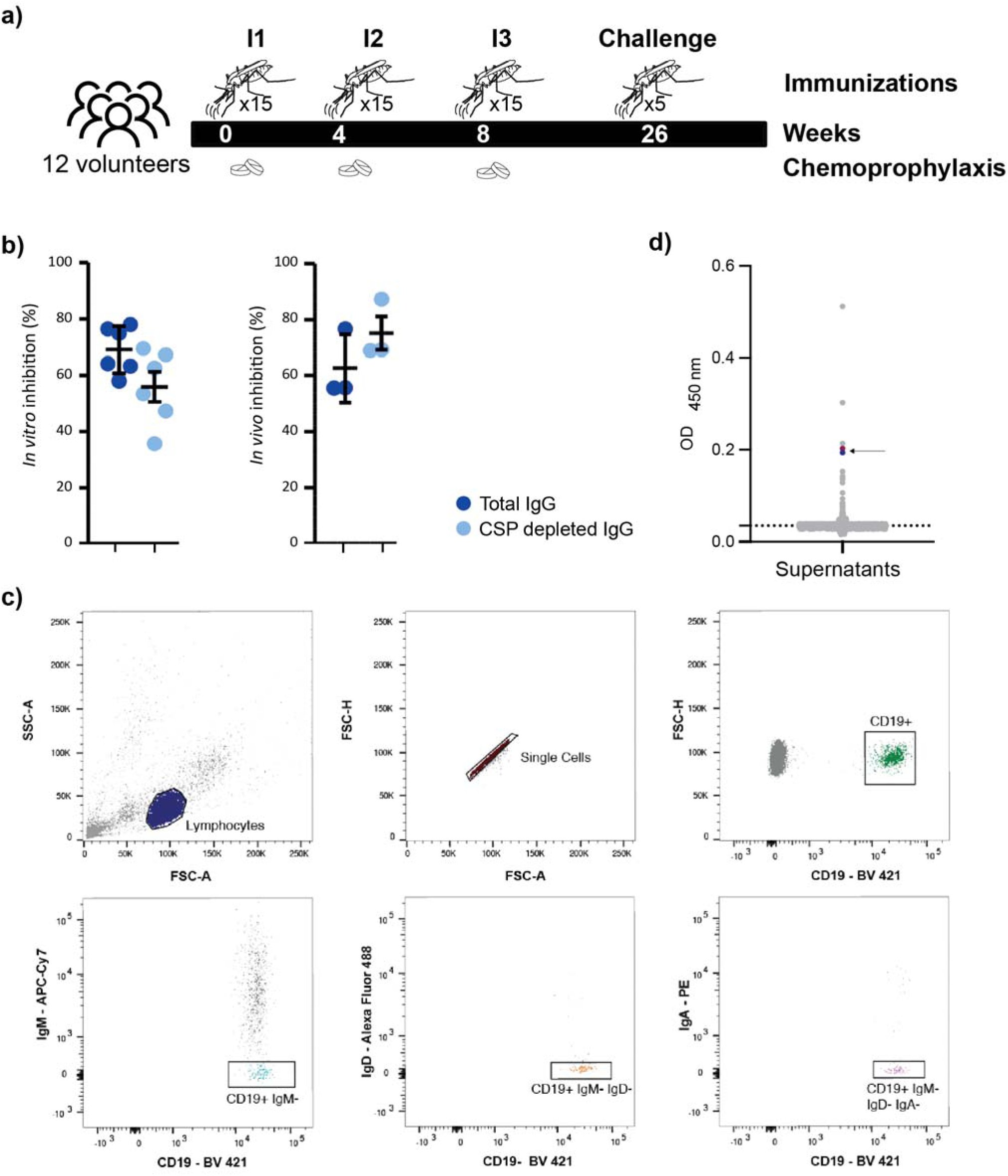
Experimental design and B-cell sorting strategy **a)** Schematic overview of CPS-based immunization trial (Walk, Reuling et al. 2017); **b)** Hepatocyte invasion inhibition of *P. falciparum* NF54 sporozoites by IgGs from volunteer 1 (Fabra-Garcia, Yang et al. 2022). Total IgG (dark blue) and IgG depleted for CSP-antibodies (light blue) were tested for invasion inhibition of the human hepatoma cell line HC-04 (left) and invasion inhibition in FRG-huHep humanized mice (right). Values represent the percentage of invasion inhibition activity. Bars represent means ± SD of one (*in vivo*) or two (*in vitro*) independent experiments, with three technical replicates each (data from (Fabra-Garcia, Yang et al. 2022)); **c)** Flow cytometry gating strategy (left to right, top to bottom) for isolation of memory B cells based on positive staining for CD19, and negative staining for IgM, IgD and IgA. **d)** Sporozoite lysate ELISA results of screened supernatants after 2-week stimulation of single sorted B cells. mAbs 16O7 and 4G16 are in pink and blue, respectively, highlighted with an arrow.

**Supplementary Figure 2:**
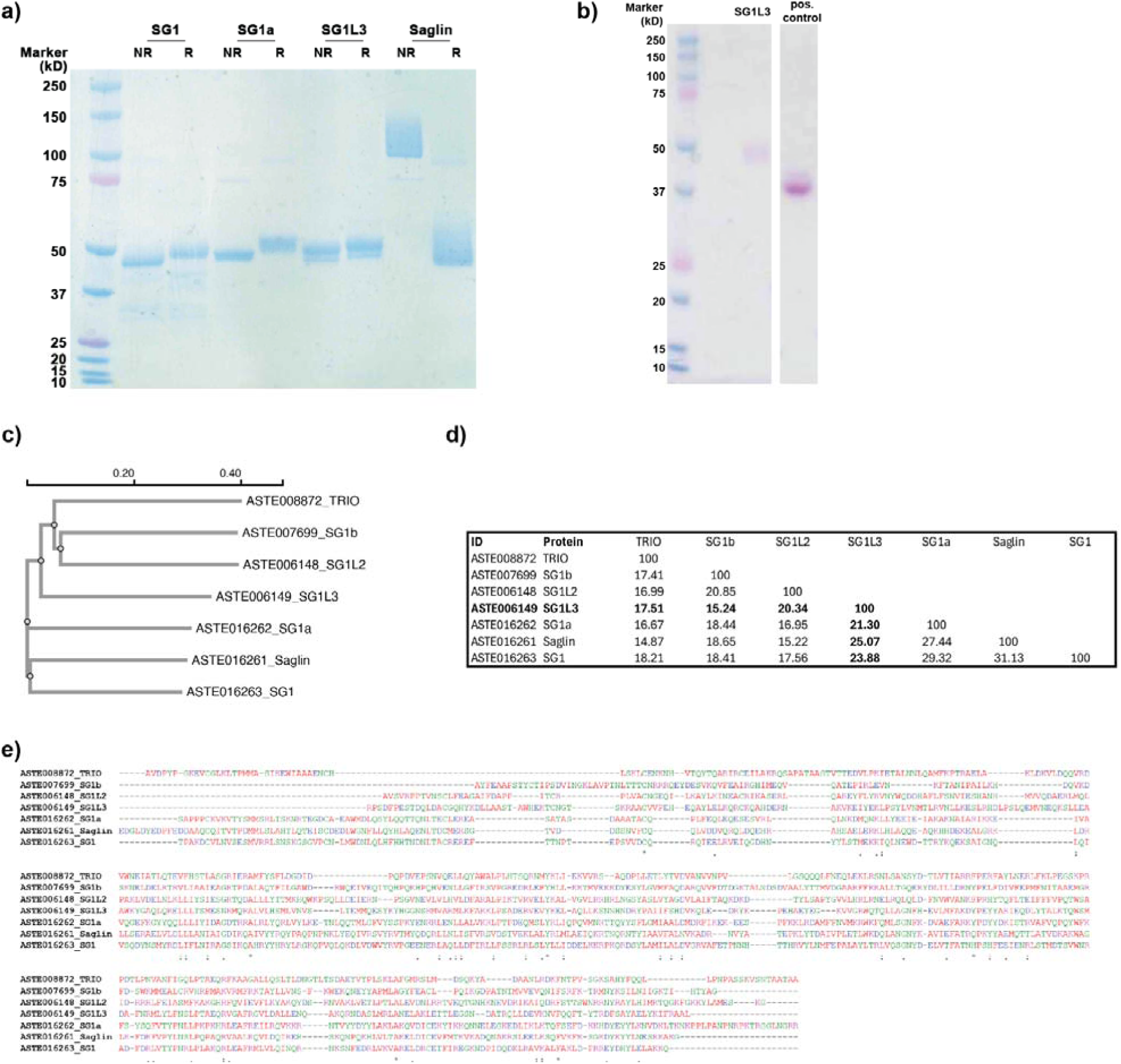
SG1 family proteins share low identity and no binding of 16O7 mAb nor SG1L3 salivary protein to gametes. **a)** Coomassie staining of recombinant SG1 family proteins under reduced (R) and non-reduced (NR) conditions. **b)** Glycostain of recombinant SG1L3 protein. Pink band indicates positive glycosylation. **c)** Phylogenetic tree of all seven SG1 family proteins from *Anopheles stephensi*; **d)** Identity matrix showing the percentage of shared identity between the SG1 family proteins; **e)** Alignment of SG1 family proteins. *(asterisk) identical,: (colon) strong, or. (period) weak degrees of amino acid conservation across sequences. The native signal peptides were omitted from the sequences; **c-e**) Generated with EMBL-EBI Job Dispatcher sequence analysis tools framework (Madeira, Madhusoodanan et al. 2024).

**Supplementary Figure 3:**
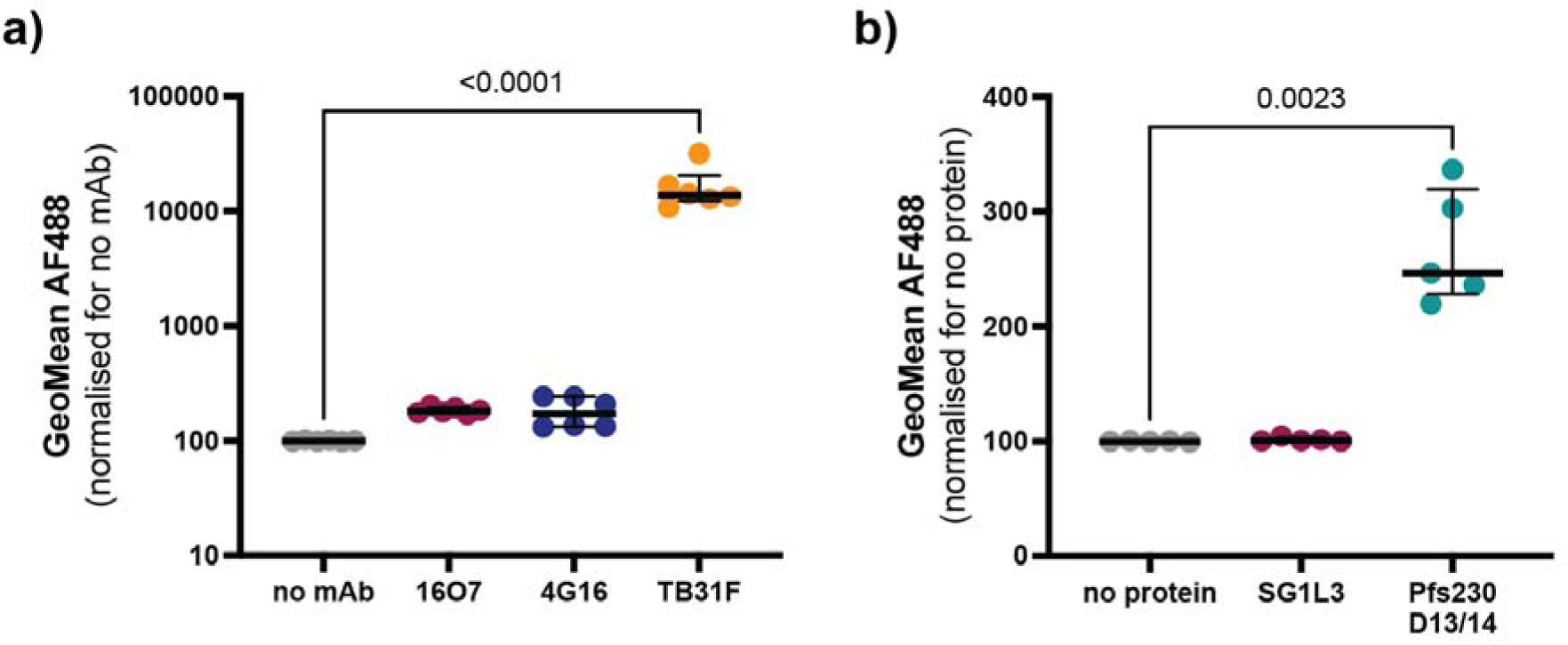
Gamete binding by monoclonal antibodies and recombinant SG1L3 **a)** Gamete binding assay by flow cytometry with different monoclonal antibodies (mAbs). Data are from two independent experiments with three technical replicates each and were normalised to the no mAb control. **b)** Gamete binding assay by flow cytometry with different recombinant proteins. Data are from two independent experiments with two or three technical replicates each and were normalised to no protein control. **a-b)** One-way ANOVA with Dunnet’s multiple comparisons test compared to no mAb/ no protein control.

**Supplementary Figure 4:**
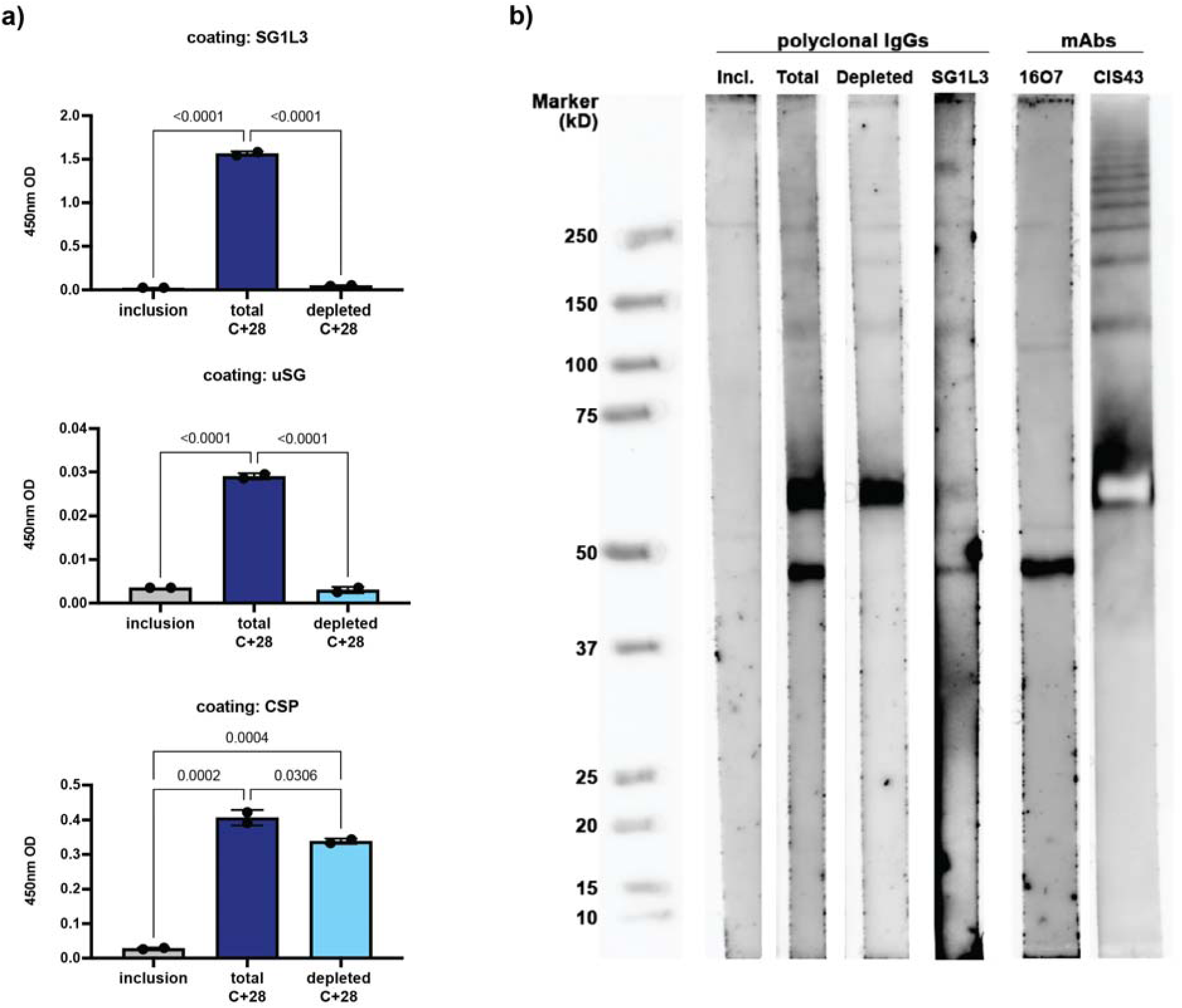
Depletion of SG1L3-specific IgGs from human volunteer IgGs **a)** ELISAs against recombinant SG1L3, uninfected salivary glands (uSG) or recombinant CSP, with inclusion, C+28 total or C+28 SG1L3-specific depleted IgGs. IgGs were tested at 15 µg/mL. All groups were compared to each other using one-way ANOVA with Tukey’s multiple comparisons correction; **b)** Western blot with NF54 sporozoite lysate, with donor polyclonal IgGs (100 µg/ml), 16O7 and CIS43 mAbs (1-2 µg/ml). Of note, purified SG1L3 condition was exposed for longer than the other conditions tested.

**Supplementary Figure 5:**
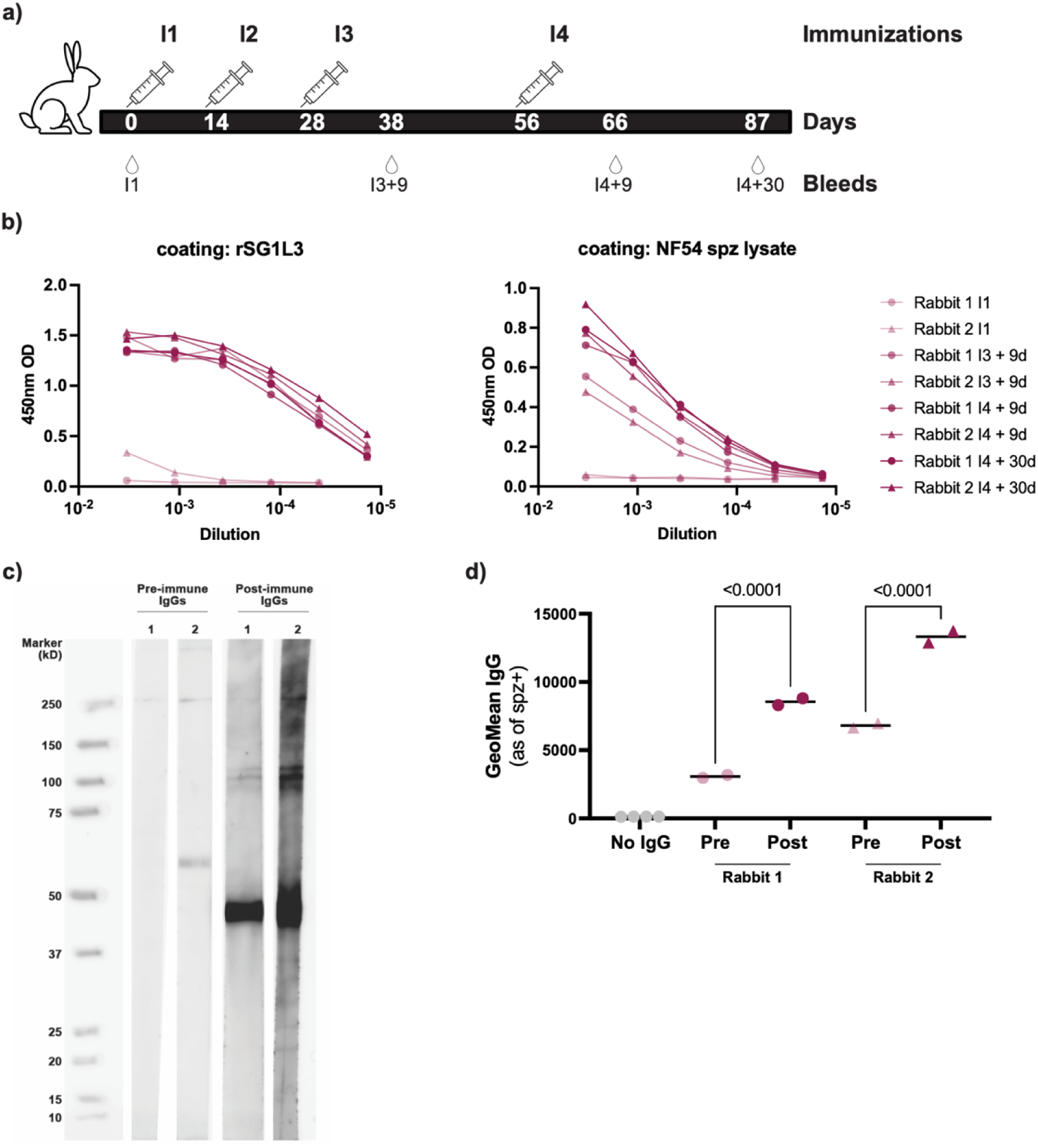
Generation of rabbit antibodies against recombinant SG1L3 **a)** Rabbit immunization scheme with the representative bleeds that were further used in experiments; **b)** ELISA with rabbit plasmas, starting at 1:300 dilution, with the different time-points pre (I1) and post-immunizations. Left, coating with recombinant SG1L3 protein; Right, coating with NF54 sporozoite lysate; **c)** Western blot with NF54 sporozoite lysate, with rabbit polyclonal IgGs (100 µg/ml), pre-immune (I1) and post-immune (I4+30), from two different rabbits (1 and 2); **d)** Post immune IgGs (10 µg/ml) from immunized rabbits recognize *Plasmodium falciparum* sporozoites by flow cytometry. Data are from one experiment with technical duplicates. One-way ANOVA with Šídák’s multiple comparisons test to each pre-immunization control.

**Supplementary Table 1:**
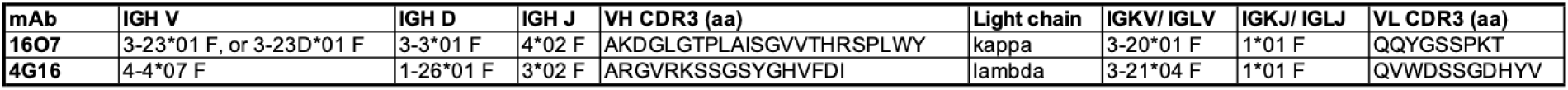
V, D, and J gene information and CDR3 sequence for 16O7 and 4G16 mAbs.

| mAb | IGH V | IGH D | IGH J | VH CDR3 (aa) | Light chain | IGKV/IGLV | IGKJ/IGLJ | VL CDR3 (aa) |
| --- | --- | --- | --- | --- | --- | --- | --- | --- |
| **16O7** | 3-23*01 F, or 3-23D*01 F | 3-3*01 F | 4*02 F | AKDGLGTPLAISGVVTHRSPLWY | kappa | 3-20*01 F | 1*01 F | QQYGSSPKT |
| **4G16** | 4-4*07 F | 1-26*01 F | 3*02 F | ARGVRKSSGSYGHVFDI | lambda | 3-21*04 F | 1*01 F | QVWDSSGDHYV |

**Supplementary Table 2:** Highest mass-spectrometry hits from co-immunoprecipitation experiment with mAb 16O7. Salivary proteins are highlighted in bold. Protein names were retrieved from *Anopheles* sialome (Arca, Lombardo et al. 2017). Intensity values are normalized mass spectrometry intensity values, searched with a 1% false-discovery rate (FDR). Raw data and full table with hits can be found in supplementary data. Order based on intensity sorted from high to low.

| Order | Intensity | Gene ID | Protein name | *A. gambiae* ID |
| --- | --- | --- | --- | --- |
| 1 | 3.06E+09 | ASTE006149 | **SG1L3** | AGAP000607 |
| 2 | 4.01E+08 | ASTE008872 | **TRIO** | AGAP001374 |
| 3 | 3.16E+08 | ASTE006148 | **SG1L2** | AGAP000609 |
| 4 | 2.63E+08 | ASTE010013 | unspecified product |  |
| 5 | 2.62E+08 | ASTE007699 | **SG1b** | AGAP000548 |
| 6 | 1.48E+08 | ASTE016224 | **SG7-2** | AGAP008215 |
| 7 | 8.61E+07 | ASTE015933 | Cytoplasmic actin | AGAP011515 |
| 8 | 8.19E+07 | ASTE009252 | Histone H4 |  |
| 9 | 7.89E+07 | ASTE016262 | **SG1a** | AGAP000611 |
| 10 | 3.42E+07 | ASTE009251 | Histone H3 |  |
| 11 | 3.37E+07 | ASTE010393 | unspecified product |  |
| 12 | 2.96E+07 | ASTE016261 | **Saglin** | AGAP000610 |

## Notes

### Competing Interest Statement

The authors have declared no competing interest.

